# Investigating the consequences of the mating system for drug resistance evolution in *C. elegans*

**DOI:** 10.1101/2024.11.07.620884

**Authors:** B. Trubenová, J. Hellinga, J. Krücken, G. von Samson-Himmelstjerna, H. Schulenburg, R. Regoes

## Abstract

The rise of anthelmintic-resistant strains in livestock threatens both animal and human health. Understanding the factors influencing anthelmintic resistance is crucial to mitigate the threat posed by these parasites. Due to difficulties in studying parasitic worms in the laboratory, the non-parasitic nematode *Caenorhabditis elegans* is used as a model organism to investigate anthelmintic resistance evolution. However, the suitability of this free-living nematode as a model for parasitic worms is debatable due to its rare androdioecious reproductive system, raising questions about the generalizability of findings from evolutionary experiments in *C. elegans* to other species. In this study, we developed a polygenic, population genetic model combined with pharmacodynamic approaches to investigate the effects of reproductive strategy and other aspects, such as dominance, mutational effects, the number of loci, and population size, on determining the dynamics and outcome of evolutionary processes. We found that androdioecious populations showed both rapid initial adaptation typical for hermaphrodites and tolerance to high drug concentrations observed in dioecious populations. They also exhibited the highest diversity and shortest time for the fixation of the beneficial allele. These results suggest that androdioecious populations can harness the advantages of both selfing and outcrossing, optimizing their reproductive strategy in response to drug selection.

## 1 Introduction

Helminths, particularly parasitic nematodes, are diverse parasites of plants, animals, and humans. Despite advances in sanitation, helminth infections are among the most common infections worldwide, affecting large human populations, mainly in low-income countries and in the poorest and most deprived communities. They cause a significant burden on the population, drastically decreasing the quality of life and life expectancy, especially when combined with malnutrition. Nevertheless, many helminthiases belong to the so-called Neglected Tropical Diseases group because they are almost absent from the global health agenda and research into these diseases is severely underfunded.

Anthelmintic drugs are generally very efficient, clearing infections within days. However, extensive use of these drugs in grazing animals led to the evolution of drug-resistant strains [Schwab et al., 2006, Schulz et al., 2018, Charlier et al., 2022, Mohammedsalih et al., 2024]. Their prevalence has increased globally in recent years, representing a severe problem in various livestock sectors [Kotze and Prichard, 2016, Sangster et al., 2018], and becoming an increasingly severe threat to agriculture and human health [Charlier et al., 2022, Wolstenholme et al., 2004, Von Samson-Himmelstjerna, 2012, Tinkler, 2020]. Indeed, the reduced sensitivity of *Ascaris* populations to benzimidazole has already been detected in children in Rwanda [Krücken et al., 2017]. As many parasitic worms are zoonotic, anthelmintic resistance in worms is a significant focus of One Health initiatives.

Understanding the factors that influence the evolution of anthelmintic resistance in parasitic worms is essential to mitigate their threat. Unfortunately, conducting laboratory experiments involving parasitic worms poses challenges due to the requirement of an experimentally accessible host organism, the lack of genetic tools for analysis of the parasites, and the long life cycle of many parasites. Consequently, these studies are relatively rare. Therefore, the intensively studied nematode *Caenorhabditis elegans* serves as a model organism for parasitic nematode infections despite not being a parasite, mainly due to its short life span and easy maintenance in the laboratory setting, well-annotated genome and similar response to anthelmintics [Bürglin et al., 1998, Geary and Thompson, 2001]. It has been used successfully as a model for parasitic worms in the past, yielding important insights into the evolution and genetics of resistance against anthelminthic drugs [Driscoll et al., 1989, Dent et al., 2000, James and Davey, 2009, Ménez et al., 2016, 2019, Hellinga et al., 2024].

However, the suitability of the free-living *C. elegans* as a model organism for parasitic worms has also been challenged [Geary and Thompson, 2001]. Beyond its free-living lifestyle, one of the key distinctions lies in its reproductive mode. *C. elegans* is androdioecious, with populations consisting of self-fertilizing hermaphrodites and a small proportion of males. While nematode reproductive modes are remarkably diverse [Singaravelu and Singson, 2011], most of the parasitic worms reproduce by mating between males and females (dioecy). One of the few known exceptions is *Phasmarhabditis hermaphrodita*, a nematode parasite of slugs, that is a hermaphrodite [Wilson et al., 1993, Rae et al., 2010]. Overall, androdioecy is relatively rare, with only a few examples [Weeks, 2012, Kanzaki et al., 2013].

This diversity in reproductive strategies raises crucial questions for understanding adaptation. Theoretical studies in population genetics have emphasized the potential impact of reproductive mode on the rate and trajectory of evolution [Orr and Otto, 1994, Kondrashov, 1994, Otto and Lenormand, 2002, Kleiman and Tannenbaum, 2009, Gerstein et al., 2011, Colegrave, 2012, Hartfield and Gĺemin, 2016, Hartfield et al., 2017, Uecker, 2017]. However, these studies often focus on the broad divide between sexual and asexual reproduction [Melían et al., 2012], rather than the finer distinctions among mating systems like dioecy, hermaphroditism, and the complex dynamics of androdioecy. As a rare exception, Husse et al. [2013] developed a one-locus model to explore the dynamics of androdioecious species, finding that a slight reproductive advantage of males can maintain high male frequencies. Weeks et al. [2006] conducted a comparative analysis of androdioecy in animals, suggesting that it may have evolved as a response to reproductive assurance in species with episodic low densities.

Due to the limited number of studies on androdioecy in animals (see Yamaguchi and Iwasa [2021] for exceptions), our understanding of the androdioecious systems and their consequences on evolutionary adaptation come primarily from plant research, where androdioecy is more common. Androdioecy requires specific conditions to evolve, and its origin and presence are still puzzling scientists [Wegewitz et al., 2008, Chasnov, 2013]. Some researchers suggest that androdioecy is an intermediate state between hermaphroditism and dioecy (see Weeks [2012] for a detailed review). It has been shown that selfing by hermaphrodites provides reproductive assurance, allowing populations to persist through periods of low density when mates are scarce [Chasnov, 2013]. In addition, selfing increases homozygosity, exposing new mutations (both beneficial and deleterious) to natural selection, thereby increasing its efficacy [Ellis and Lin, 2014]. However, selfing may reduce genetic variation over time, limiting the potential to adapt to novel conditions. In contrast, when hermaphrodites outcross with males, new genetic variation can be introduced into the population, which may be especially suitable for adaptation to changing environments [Yamaguchi and Iwasa, 2021, Ellis and Lin, 2014]. Furthermore, outcrossing is expected to compensate for lower fitness caused by inbreeding depression [Frankham et al., 2010].

The evolution of androdioecy in *C. elegans* has already been studied by experimental and modelling approaches, taking into consideration mating ability and spontaneous production of males [Chasnov and Chow, 2002]. Chasnov [2010] suggests that dioecy can transition to androdioecy when the benefits of reproductive assurance outweigh the costs of selfing. Using experimental evolution, Anderson et al. [2010] show that males can be maintained at high frequencies when populations have genetic diversity or are exposed to novel environments. Teotonio et al. [2012] suggest that in androdioecious populations with standing genetic diversity, outcrossing rates were maintained at an optimal level of around 50% through stabilizing selection on male frequencies. On the other hand, according to Stewart [2003], males appear to reproduce at a rate just below that necessary for them to be maintained. However, the previous models usually assumed incomplete mating abilities of the males, which is particularly true for the lab-adapted *C. elegans* strain N2, but not for most natural isolates of *C. elegans* [Wegewitz et al., 2008, Teotónio et al., 2006]. Therefore, Cutter et al. [2019] state that males in *Caenorhabditis* are understudied compared to hermaphrodites while playing an important role in sexual selection, sexual conflict, and reproductive mode transitions. In summary, the androdioecious reproductive mode presents a complex landscape of potential evolutionary advantages and disadvantages. However, due to the limited number of studies, the evolutionary consequences of androdioecy remain poorly understood and the generalizability of findings from *C. elegans* to other nematode species with diverse reproductive modes remains an open question.

To address this gap, this study investigates the suitability of *C. elegans* as a model organism for studying the evolution of antihelmintic resistance in parasitic nematodes. We aim to improve our understanding of the evolutionary consequences of androdioecy using a mathematical modelling approach. For this, we developed a polygenic population genetic model of a nematode population with dioecious (mating), hermaphroditic (selfing), or androdioecious (combined selfing and mating) reproduction. We investigate how these reproductive modes influence the rate and outcome of adaptation to an anthelmintic-containing environment. Moreover, we investigate how other factors, such as dominance, population size, the number of loci, and the size of their effects influence the consequences of the mating system. To incorporate the selective pressures of this environment, we utilize a modified pharmacodynamic framework from Regoes et al. [2004], which captures the varying effects of drug concentrations on populations with different mutations, thus enabling us to model anthelmintic resistance in diploid organisms. This approach, combined with the population genetics [Trubenová et al., 2022], promises novel insights into the role of reproductive strategy in drug resistance evolution among parasitic worms.

## 2 The model

Below, we describe our compartmental, polygenic, population genetic model. Inputs of the model are various parameters determining the genetics (e.g., number of loci), the translation to phenotype (e.g. fitness effects of individual mutations), and the details of the experiment (e.g. length of the treatment, drug concentration), see Table 1 for details. The model compartments represent a particular class of individuals defined by their sex and genotype. We assume that the organisms are diploid.

**Table 1:** Symbols used in the model.

| Category | Variable | Explanation | Variable type |
| --- | --- | --- | --- |
| population | N | Initial number of worms in the simulated population | input |
| Genetics | k | Number of loci | input |
| | $\mu$ | Mutation rate per genome (not locus) | input |
|  | **g** | Genotype, vector of length k with values 0, 1 or 2 | calculated |
|  | **d** | Dominance, vector of length k with values 0 or 1 | input |
|  | **p** | Phenotype, vector of length k with values 0 or 1 | calculated |
|  | **b** | Benefit vector, describing benefits associated with mutation at each locus | input |
|  | **c** | Cost vector, describing cost associated with mutation at each locus | input |
| Pharmacodynamics | C | Total cost associated with a genotype | calculated |
|  | B | Total benefit associated with a genotype | calculated |
|  | A | Drug concentration | input |
| | $\kappa$ | Hill's coefficient describing steepness of the curve | input |
| | $EC_{50}$ | Drug concentration that produces 50% of its maximum possible effect | input/calculated |
| | $\phi$ | Fitness | calculated |
| Reproduction | $f_m$ | Fraction of males | calculated |
| | $\xi$ | Fraction of hermaphrodites that mate | input |
| | $egg_s$ | Mean selfing fecundity/ offspring number | input |
| | $egg_m$ | Mean mating fecundity/ offspring number | input |

How each genotype manifests depends on whether the mutations are dominant or recessive. Each genotype is associated with a basal fitness (in the absence of the drug), and it has an *EC*_50_ value (concentration of a substance that produces 50% of its maximum possible effect), as explained in detail below. Hermaphrodites from each compartment can self-fertilize or mate, laying a number of eggs according to the origin of sperm, their fitness, and genotype. These eggs may have various genotypes, assuming that genetic loci are not linked and are carried at different chromosomes. Additionally, mutations may occur. Finally, offspring generation replaces the parental one and may be reduced to a required size depending on the modelled scenario. All of these processes are explained in detail below. See section Simulation process for individual steps and Figure 1 for a schematic representation of the overall simulation process.

**Figure 1:**
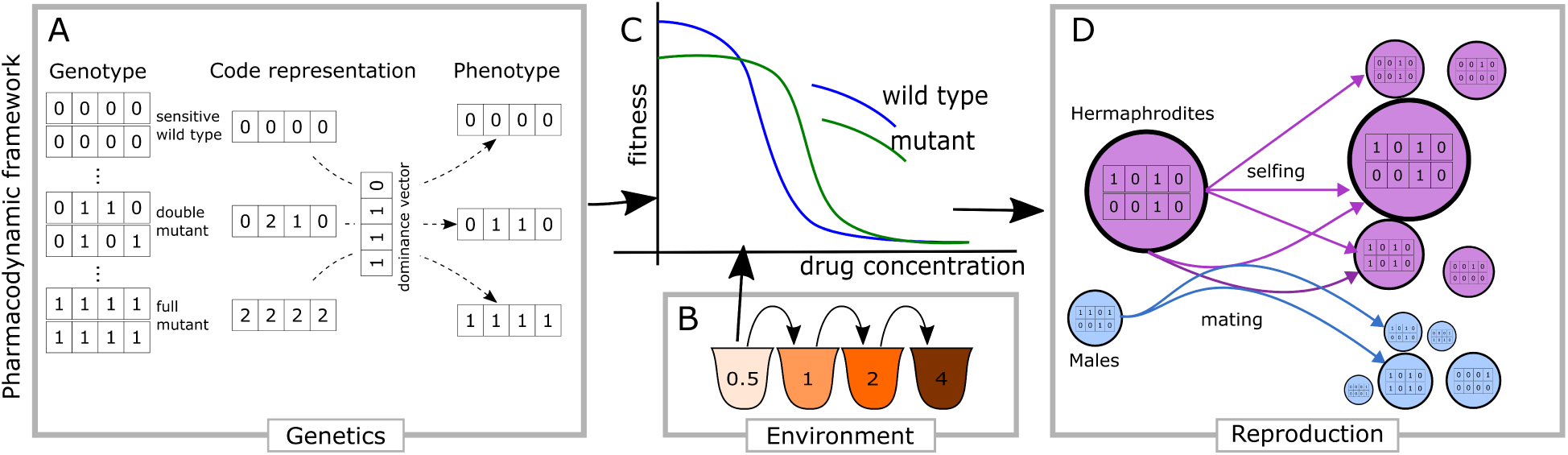
Modelling evolution of resistance in diploid organisms. Panel A shows diploid genomes with four loci, each with two alleles (0 or 1), whereby 1 represents a mutation. Panel B represents a serial passage experiment with increasing drug concentration. Panel C shows drug resistance for wild-type and mutant strains, indicated through the relationship between drug concentration (X axis) and fitness (Y axis). Panel D highlights different reproduction alternatives between a female/hermaphrodite (in purple) and a male (blue). While selfing can generate offspring of various genotypes, all are hermaphrodites. Mating produces various genotypes equally split between males and hermaphrodites.

The model’s output is a complete record of the population sizes of all compartments over time. Therefore, it is possible to observe population dynamics (e.g., changes in population sizes, allele frequencies, or accumulation of the mutations), the timing of events (e.g., when a mutant appears or reaches a certain frequency), and to calculate the emerging population properties (e.g., diversity) and observe their changes over time. The model is implemented in Python. The simulations are stochastic, allowing us to determine the probability distributions of the outcomes.

### Genotypes

Parasitic worms, as well as *C. elegans*, are diploid organisms, in which the drug resistance is known to be encoded by multiple loci [Dent et al., 2000, Ménez et al., 2016, Von Samson-Himmelstjerna et al., 2007, Page, 2018, Mcintyre et al., 2025]. Therefore, we represent their genotype, consisting of *k* unlinked loci relevant for the modelled adaptation process, as a string of length *k*. Each position describes the number of mutated alleles at that particular locus: 0 represents a sensitive homozygote (at the particular locus), 1 represents a heterozygote, and 2 represents a homozygote with two mutated alleles. With *k* loci, there are 3*^k^* possible genotypes. These represent sub-populations (classes) with possibly distinct pharmacodynamic properties.

As we assume a diploid genome, the relationship between the alleles of the same locus must also be defined. Vector **d** (length *k*) determines the dominance of mutation at individual loci. If *d_j_* = 1, the mutated allele at locus *j* is dominant, and genotypes that have one or two mutated alleles at this locus manifest in the same way. If *d_j_* = 0, the mutated allele at locus *j* is recessive, and only homozygous mutations lead to a fitness effect. Using **d**, for each genotype **g** we determine phenotype **p** (length *k*), with its elements capturing whether the effect of each mutated locus should be considered or not (see Pharmacodynamics and fitness).

### Sexes

From each genotype, it is possible to have individuals who are capable of contributing their genetic information to the next generation but do not lay eggs, further referred to as males. In addition, we also include individuals who contribute their genetic information and are capable of laying eggs. Depending on the simulated scenario, these individuals may be able to self (hermaphrodites) or require mating in order to lay eggs (representing females). However, due to simplicity, we will refer to all egg-laying individuals as hermaphrodites throughout this manuscript, and the ability to lay eggs with or without male presence will be modulated by a parameter *ξ*, as explained below. Individuals of both sexes may have any possible genotype, altogether creating 2 *×* 3*^k^* of sub-populations (compartments) that are simulated.

### Pharmacodynamics and fitness

The effects of anthelmintic drugs on worms can vary at different stages of their life cycle. These drugs can kill adult worms, paralyze them, prevent them from reproducing, or block the hatching of their eggs. In our model, we simplified the impact of anthelmintics on worm fitness by focusing solely on a reduced number of hatched eggs and thus the number of offspring a particular genotype can produce, regardless of the underlying causes. Moreover, we define fitness as the number of offspring that survive until adulthood and are able to reproduce. In this sense, fitness is a result of various processes - from decreased survival of the genotype in focus, with subsequently reduced offspring production and decreased mobility, thus affecting the ability to feed/find a mating partner(s), or just having fewer eggs per individual.

In the presence of an anthelmintic, the fitness of all individuals is reduced. The concentration of the anthelmintic required to produce a specific effect (such as killing, paralyzing the worms, or preventing reproduction) is higher for the mutated worms, indicating an increase in the *EC*_50_ compared to the sensitive strain. However, we also assume that mutations conferring drug resistance come with a cost in the absence of the drug, leading to a reduction in fitness.

Therefore, in our model, each resistant allele is associated with a benefit increasing the carrier’s *EC*_50_ in the presence of drugs and a cost-reducing fitness in its absence. Costs and benefits are given as vectors **c** and **b** of length *k*, respectively, with each element corresponding to the respective locus. Mutational effects of resistance-conferring mutations can be of various sizes and can interact together [Das et al., 2020]. To avoid introducing epistatic effects, we assume that both costs and benefits combine in an additive manner [Trubenová et al., 2022, Knopp and Andersson, 2018, Sackman and Rokyta, 2018, Wistrand-Yuen et al., 2018, Igler et al., 2021].

The total cost *C_i_*associated with phenotype *i* is

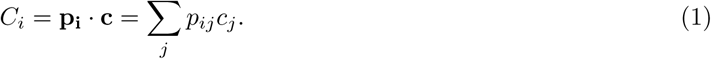

and the total benefit *B_i_* associated with phenotype *i* is:

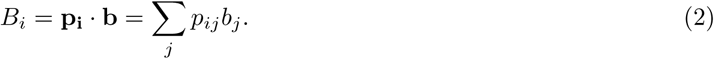

The total cost associated with a particular genotype (strain) determines its fitness *ϕ_i_* in the absence of the drug *ϕ_i_*(0) = 1 *− C_i_*, where the fitness of the fully sensitive strain is 1. The total benefit captures the increase of the *EC*_50_ of this strain as *EC*_50_*_i_* = *EC*_50_*_wt_* + *B_i_*, where *EC*_50_*_wt_* is the *EC*_50_ of the sensitive wild-type. The fitness in the absence of the drug and *EC*_50_ define pharmacodynamic curves that allow us to determine the expected fitness of any strain *i* (see Figure 2):

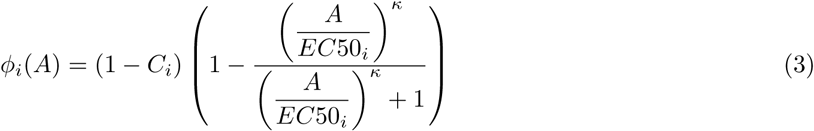

**Figure 2:**
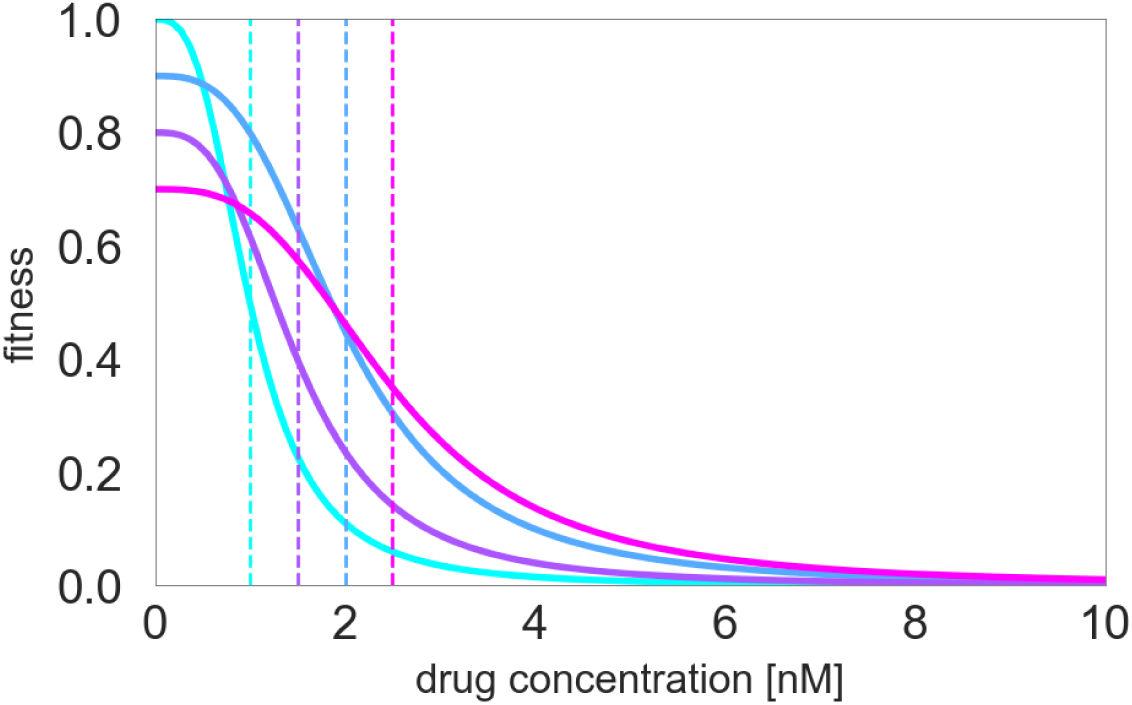
Fitness function of different strains. In this illustrated example, resistance is encoded by two independent recessive mutations. Sensitive strain (cyan) has the highest fitness in the absence of the drug but the lowest *EC*_50_ (shown as a dashed line). Two different single mutants (with homozygous mutations at one of the two loci, shown in blue and purple) have lower fitness but higher *EC*_50_. The double mutant (homozygous for mutations at both loci, shown in purple) has the lowest fitness of all the strains in the absence of the drug but the highest *EC*_50_. Mutations are recessive, thus, heterozygous loci do not contribute to resistance, nor do they reduce fitness.

where *A* is the drug concentration and *κ* defines the steepness of the curve.

### Egg count

Apart from fitness, the number of eggs the hermaphrodite can lay is also influenced by the origin of sperm: whether it comes from mating or selfing. In line with the experimental observations in *C. elegans*, we will assume that mated, wild-type individuals can lay, on average, 1000 eggs in a drug-free environment when out-crossed (*egg_m_* = 1000) but only 300 when selfing (*egg_s_* = 300) [Kimble and Ward, 1988]. Therefore, the number of eggs that the individual with genotype *i* lays is 300*ϕ_i_*(*A*) if the individual reproduces by selfing, or 1000*ϕ_i_*(*A*) if outcrossed.

### Reproduction

The fraction of males in the population is denoted *f_m_*, the rest are hermaphrodites. Depending on the simulated scenario, a fraction *ξ* of hermaphrodites in the population engage in mating with males, while the remainder reproduces through self-fertilization. Mating between hermaphrodites and males is random with respect to male genotype and male fraction *f_m_*, provided that there are at least some males. Thus, the frequency of mating between a hermaphrodite and a male of a given genotype is proportional to the frequency of that male genotype in the population. Offspring are generated with probabilities calculated using Punnet squares. Half of the offspring (see the definition of egg count) in each genotype class are males, while the other half are hermaphrodites, merged with the populations calculated in the selfing step outlined below.

Due to recombination and/or chromosomal shuffling, selfing individuals can also produce offspring with different genotypes, but these are always hermaphrodites. As the formation of males during selfing is rare, in our model, we assume that males are only generated by mating. A probability of a hermaphrodite that belongs to the genotype *i* to generate eggs/offspring that belongs to the sub-population with genotype *j* is calculated based on the Mendelian rules using Punnet squares for *k* loci. The linkage between loci is not currently considered; any allele can be inherited independently from another allele at a different locus.

### Mutations

During reproduction, mutations may occur. We assume that one mutation in the genome occurs directly after fertilization (either by mating or selfing) with probability *µ*, changing one locus in the whole genome from 0 to 1, 1 to 0 or 2, or from 2 to 1. Only a single mutation is allowed per individual, so approximately (sampling from Poisson distribution) *µN_i_* individuals from genotype class *i* can mutate into any genotype class in the mutational distance 1. If there are multiple target genotype classes, *µN_i_* individuals are divided into all of them in equal proportions.

### Environment

The experiment, or the treatment, consists of any number of cycles (length of the cycle corresponds to a generation time), starting drug concentration, changes in drug concentrations and their schedule, initial population sizes, the drug degradation rate, conditions for the environmental change (e.g. increase of concentration), and the dilution of the final population before it enters the next cycle. By setting these parameters, many different treatment regimes can be modelled: from a single treatment through serial passages commonly used in evolutionary experiments to a chemostat environment.

### Simulation process

First, the starting population of the desired size is seeded in the wild-type compartment, split between hermaphrodites and males according to the simulated scenario (see section Simulated scenarios). The initial drug concentration is set, and a defined number of cycles corresponding to generations is simulated according to the following procedure:

1. The fitness of all strains is calculated based on the defined environment (drug concentration) using their respective pharmacodynamic curves (Figure 2).
2. A given fraction of hermaphrodites (*ξ*) mates with males, laying a number of eggs according to the fitness of the strain they belong to. The newly generated offspring are allocated to various genotypes based on the mated pair (see above) and split into hermaphrodites and male compartments in a ratio of 1:1.
3. The remaining hermaphrodites reproduce by selfing, laying a number of eggs according to the fitness of the strain they belong to. These offspring are allocated to various compartments based on the generated genotypes (see above). All are assumed to be hermaphrodites.
4. The fraction of offspring is calculated from the mutation rate and population size; these offspring mutate and are then allocated to different compartments (see above).
5. The offspring population replaces the parental population.
6. If the population increases at least 20-fold, the population is deemed sufficiently adapted and diluted to the original size. The concentration is altered based on a pre-defined rule given by a studied scenario (e.g. kept constant or increased according to a given sequence).
7. If the population does not increase at least 20-fold, it is not considered sufficiently adapted, and the drug concentration remains unchanged. Nevertheless, the population is diluted to the original size if necessary.
8. If the population goes extinct, the simulation is terminated.

## 3 Simulated scenarios

We take advantage of this study’s computational nature and investigate a wide range of parameters and aspects that would not be feasible in a laboratory setting. For each scenario, we conducted 100 repeats. Moreover, we followed the populations for 500 generations, which would correspond to more than 4 years of continuous evolution experiments with *C. elegans*. Below, we explain the different aspects we focused on and the parameters that we used.

### 3.1 Mating systems

In our simulations, the fraction of hermaphrodites that mate (*ξ*) is independent of the fraction of males in the population. This corresponds to a biological scenario where the availability of males is not a limiting factor, but rather, the mating fraction is determined by the avoidance or inclination of hermaphrodites to mate. This means that if any males are present in the population, the fraction of hermaphrodites *ξ* will mate, with the frequency of matings contributed by males of different genotypes proportional to the frequency of those genotypes in the male population. By varying the parameter *ξ*, we can simulate different mating systems, from hermaphroditic through androdioecious to dioecious.

As males are only generated during mating, their fraction is directly influenced by the fraction of mating hermaphrodites:

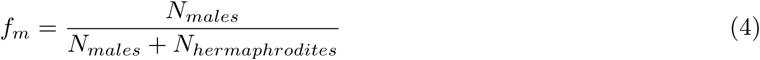

where the number of male offspring *N_males_* = 0.5*ξegg_m_N_H_* and *N_H_* is the number of hermaphrodites in the parental generation. The number of hermaphrodite offspring is *N_hermaphrodites_* = *ξegg_m_N_H_* + (1 *− ξ*)*egg_s_N_H_*, where the first term is generated by mating and the second term by selfing. Therefore, the fraction of males is

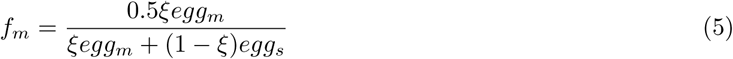

and it is independent of *N_H_*, as well as the initial fraction of males (provided there were some males). It only depends on the mating fraction of hermaphrodites *ξ*, and the fecundity of hermaphrodites when selfing and mating. For simplicity, we initialize all simulations with a fraction of males of 0.5.

To simulate the **hermaphroditic mating system**, we set the fraction of hermaphrodites that mate to *ξ* = 0. This means that males are purged from the population after a single generation, and the population remains hermaphroditic.

On the other hand, to simulate the **dioecious mating system**, we set the fraction of mating hermaphrodites to *ξ* = 1. This means that all the egg-laying individuals reproduced only by mating, essentially representing females. Due to the stochasticity of the system, the fraction of males fluctuates around *f_m_* = 0.5.

When 0 *< ξ <* 1, some hermaphrodites reproduce by mating, while others by self-fertilization, representing an **androdioecious mating system**. We simulated a range of values of *ξ* in increments of 0.1, where 0.1 *≤ ξ ≤* 0.9. The fraction of males fluctuates around *f_m_* given by equation 5.

### 3.2 Genetic factors

#### Number of loci

First, we assume two loci with relatively small effects (cost 5 %, benefit 0.5 nM), resulting in nine possible genotypes. Then, to investigate evolutionary dynamics in more complex scenarios, we assume six independent loci, three with small effects (cost 5 %, benefit 0.5 nM) and three with large effects (cost 10 %, benefit 3 nM). It allows the generation of 729 different genotypes, reflecting the complexity found in nature. Note that the fitness (in the absence of a drug) of the sensitive strain (with no mutation) is 1, and its *EC*_50_ is 1 nM. The mutation rate is *µ* = 0.0001 mutations per genome.

#### Dominance

We investigate adaptation when all loci are dominant, or all loci are recessive.

### 3.3 Size and timing

#### Population size

Size of the population is known to play an important role in the rate of adaptation. Therefore, we performed simulations with 200, 2000, 20000 and 200000 individuals.

#### Time scale

When evaluating the evolutionary consequences of the mating strategy, it is important to consider time scales. While some mating strategies may encourage short-term survival and rapid adaptation, others enable better survival in the long run. Therefore, we performed 100 simulations of 500 generations for each parameter set. Having complete information about the dynamics (population sizes of all compartments at all times), we compared observations made at different time points and discussed conclusions that would be made if only the particular time point was investigated. In particular, we looked at populations after 10, 25, 100, 250, and 500 generations.

### 3.4 Environment

Inspired by *in-vitro* evolution experiments performed in *C. elegans* to investigate drug resistance evolution, we simulated a gradually increasing drug concentration (e.g. ivermectin), with concentration increases following a pre-defined sequence shown in Table 2. The population starts in an environment with no drug concentration, which increases over time when the population is deemed sufficiently adapted (having an average egg count of at least 20 per individual). Populations are passaged from one generation to another, increasing the drug concentration but allowing constant growth (no limits imposed by carrying capacity). This approach has been shown to enable the evolution of high resistance while limiting the probability of extinction [James and Davey, 2009, Ménez et al., 2016, Hellinga et al., 2024], which may occur if the concentration is increased too steeply. Supplementary Figure S1 A shows preliminary simulations of populations exposed to 10 nM drug concentration without prior exposure to lower drug concentrations. The majority of these populations go extinct within the first 20 generations, especially in smaller populations. Moreover, gradual exposure to drugs leads to the survival of all populations regardless of their size (Supplementary Figure S1 B) and their addaptation to 10 nM concentration or even higher (Supplementary Figure S1 C).

**Table 2:** Sequence of ivermectin concentrations.

|  |  |  |  |  |  |  |  |  |  |  |  |  |  |  |  |  |  |  |  |  |  |  |  |  |
| --- | --- | --- | --- | --- | --- | --- | --- | --- | --- | --- | --- | --- | --- | --- | --- | --- | --- | --- | --- | --- | --- | --- | --- | --- |
| 0 | 0.1 | 0.2 | 0.4 | 0.8 | 1 | 1.5 | 2 | 2.5 | 3 | 4 | 5 | 6 | 8 | 10 | 12 | 15 | 18 | 21 | 24 | 27 | 30 | 35 | 40 | 45 |

**Table 3:** Performed simulations.

| Loci | Dominance | N | $\xi$ |
| --- | --- | --- | --- |
| 2 | 0 | 200 | 0-1 |
|  |  | 2000 | 0-1 |
|  |  | 20000 | 0-1 |
|  |  | 200000 | 0-1 |
| 2 | 1 | 200 | 0-1 |
|  |  | 2000 | 0-1 |
|  |  | 20000 | 0-1 |
|  |  | 200000 | 0-1 |
| 6 | 0 | 200 | 0-1 |
|  |  | 2000 | 0-1 |
|  |  | 20000 | 0-1 |
|  |  | 200000 | 0-1 |
| 6 | 1 | 200 | 0-1 |
|  |  | 2000 | 0-1 |
|  |  | 20000 | 0-1 |
|  |  | 200000 | 0-1 |

### 3.5 Implementation and writing

The code was developed and written in Python, performing simulations summarised in 3. Generative AI (Gemini and Grammarly) was used to improve the grammar and clarity of this article.

We performed simulations for dominant (value 1) or recessive (value 0) mutations, four population sizes, and 11 values of *ξ* (hermaphrodites, 9x androdioecious populations with an increment of 0.1, dioecious populations). All simulations were run for 500 generations, with 100 repeats. Altogether, we performed 8800 simulations.

## 4 Results

In this study, we simulated the evolution of worm populations in an environment with increasing concentrations of ivermectin as a model to study nematode resistance evolution. The concentration was raised when the population was deemed sufficiently adapted to its current conditions. Thus, the rate of increase in drug concentration acts as a proxy for the rate of adaptation. We examined how various genetic and environmental factors, along with different mating strategies, influenced the rate of adaptation and the maximum concentration that the populations reached over a specified period.

### 4.1 Mating facilitates sustained adaptation in large populations with recessive mutations

Our findings demonstrate that a gradual drug concentration increase enables populations to adapt to relatively high drug concentrations (Figure 3). Population size is a critical factor, with larger populations exhibiting, in general, faster adaptation. As simulations are initiated with wild-type populations (no mutations are present), it suggests that mutational supply limits adaptation in smaller populations, and mutations are less frequent in smaller populations.

**Figure 3:**
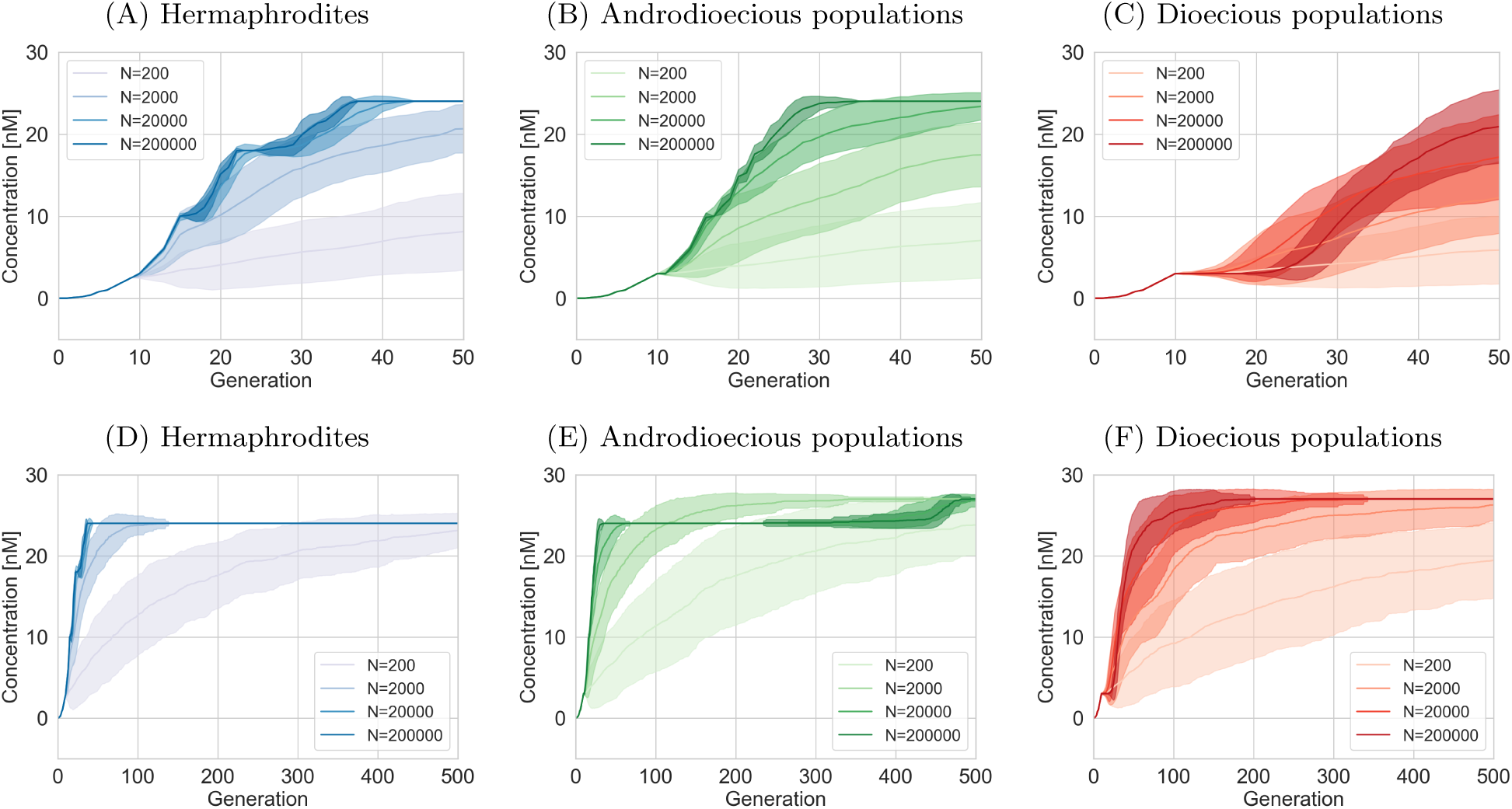
Adaptation to increasing drug concentration, assuming recessive mutations and six loci. Mean and standard deviation of 100 simulation trials. In androdioecious populations, 50% of hermaphrodites reproduce by mating and 50% by selfing. A-C) First 50 generations. D-F) All 500 generations.

However, the interplay between population size and adaptation is more complex, as is particularly evident in Figure 3 C, for mating populations. Here, larger populations (200000 individuals) initially lag behind smaller ones (2000 and 20000 individuals) in progressing to higher concentrations. A similar effect is also seen in androdioecious populations (*ξ* = 0.5), where populations of 2000 individuals are the first to overcome the 24 nM threshold (Figure 3 E). This indicates that while the mutation supply limits small populations, once the mutation appears, it takes longer for the mutant to outcompete the others in larger populations.

Furthermore, Figure 3 highlights distinct adaptation patterns across the three mating strategies. Strictly selfing populations (hermaphrodites only) rapidly progress to 24 nM but fail to exceed this threshold within the simulation timeframe (Figure 3 D). In contrast, strictly mating populations exhibit slower initial adaptation but then continue to progress to 28 nM (Figure 3 F). Androdioecious populations display a rapid initial adaptation, similar to that of hermaphrodites, followed by a slower progression to higher concentrations. Nonetheless, they ultimately adapt to a greater extent than hermaphrodites (Figure 3 E). Moreover, adaptation of the smallest androdioecious populations in the given timeframe exceeded that of the smallest dioecious populations. This suggests that androdioecious populations may be able to reap the benefits of both mating and selfing strategies. Therefore, evolution experiments with *C. elegans* might overestimate the adaptation to anthelmintics, especially at low population sizes.

### 4.2 Rate of adaptation to higher concentration does not directly reflect genetic changes

The increasing ability to survive in higher concentrations is enabled by the gradual accumulation of resistance mutations. However, the rate of adaptation does not necessarily reflect the rate of mutation accumulation, as other differences between reproductive strategies are also present.

Figure 4 illustrates the average number of homozygous loci within the population over time, considering that resistance mutations are recessive, with six loci potentially contributing to resistance. Hermaphroditic populations exhibit a faster accumulation of homozygous mutations compared to strictly mating populations, which demonstrate the slowest adaptation. Furthermore, hermaphroditic populations display a stepwise increase in homozygous loci, whereas other mating systems exhibit a more gradual progression.

**Figure 4:**
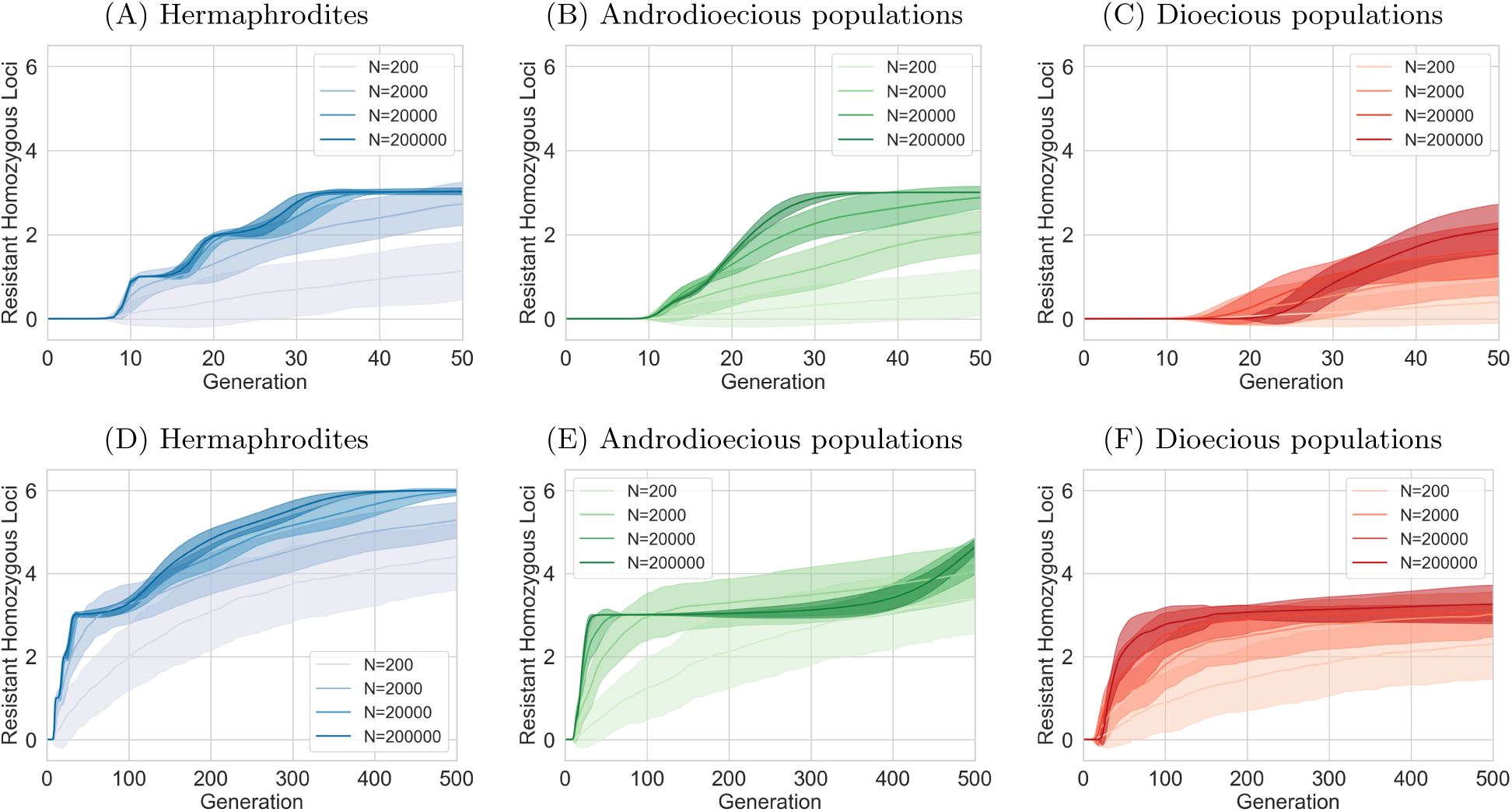
Number of homozygous loci over time, assuming recessive mutations and six loci. Mean and standard deviation of 100 simulation trials. In androdioecious populations, 50% of hermaphrodites reproduce by mating and 50% by selfing. A-C) First 50 generations. D-F) All 500 generations.

While Figure 4 shows that hermaphrodites reach the maximum of six homozygous loci within the simulated period, this is not observed in dioecious populations. Only three (or fewer) loci with large effects are homozygous for the mutated allele in strictly mating populations by the end of the simulations. Androdioecious populations display a similar pattern, remaining stagnant at three homozygous mutated loci for an extended duration. However, Figure 4 B shows that this threshold is eventually crossed, and these populations continue to accumulate mutated loci.

Initially, this appears to contradict the earlier observation of dioecious populations achieving resistance to the highest concentrations. Therefore, it is important to note here that in the same conditions, dioecious populations produce, on average, a greater number of offspring (1000 per hermaphrodite, so 500 per individual) than selfing hermaphrodites (300 per individual; see the Reproduction section) in a drug-free environment [Kimble and Ward, 1988]. This difference in average egg count persists across genotypes and drug concentrations, with hermaphrodites always laying more eggs after mating compared to self-fertilization. It allows dioecious populations to have satisfactory fertility to meet the requirements for increasing drug concentration. Consequently, the selection pressure on dioecious populations for further evolution is reduced, decelerating the accumulation of mutations. At the same time, hermaphrodites, while having all six loci adapted, do not progress to higher concentrations. This is caused by the small effects of three of the loci. Even their combined effect is not sufficient to allow them to reach the egg count threshold required to proceed to higher concentrations (see the section on Genetics and sections below).

The androdioecious populations also slow down mutation accumulation after the loci of large effect are fixed, but eventually accumulate mutation with small effects faster than dioecious populations, as selection pressure is higher. As we see in Figure 3, large populations reach some kind of adaptation threshold after approximately 30 generations. This is because three loci with large effects are already mutated, and these three mutations are now fixed in the population. The populations progressed quickly into high concentration, where they were surviving but not doing well enough to proceed into higher concentrations (not producing an average egg count of at least 20). The remaining smaller mutations are not yet fixed, which we can see in Figure 4 B (the average number of mutated loci is just three). However, these mutations with small effects are present and slowly (because of their small effect) increase in frequency. After about 400 generations, their combined effect is strong enough to allow concentrations to increase (in some of the simulations), which, in turn, increases selection pressure, and the mutation accumulation of these three remaining mutations with small effects accelerates (as seen in Figure 4 E and Figure 3 E).

These observations have implications for the relevance of evolution experiments using *C. elegans* as a model organism for dioecious parasitic worms. Our results suggest that these experiments might overestimate the number of resistance mutations that would evolve in natural parasitic populations.

### 4.3 Androdioecy accelerates fixation of beneficial mutations

If the environment changes rapidly, the beneficial allele can increase in frequency quickly. Figure 5 depicts the time to fixation (defined as reaching 95% frequency) of the first homozygous mutation across various mating systems, with the proportion of hermaphrodites engaging in mating represented by *ξ*. This ranges from exclusive selfing in hermaphrodites (*ξ* = 0, shown in blue), through androdioecious populations (0 *< ξ <* 1, green), to dioecious populations (*ξ* = 1, red). Multiple panels within the figure illustrate results for different population sizes.

**Figure 5:**
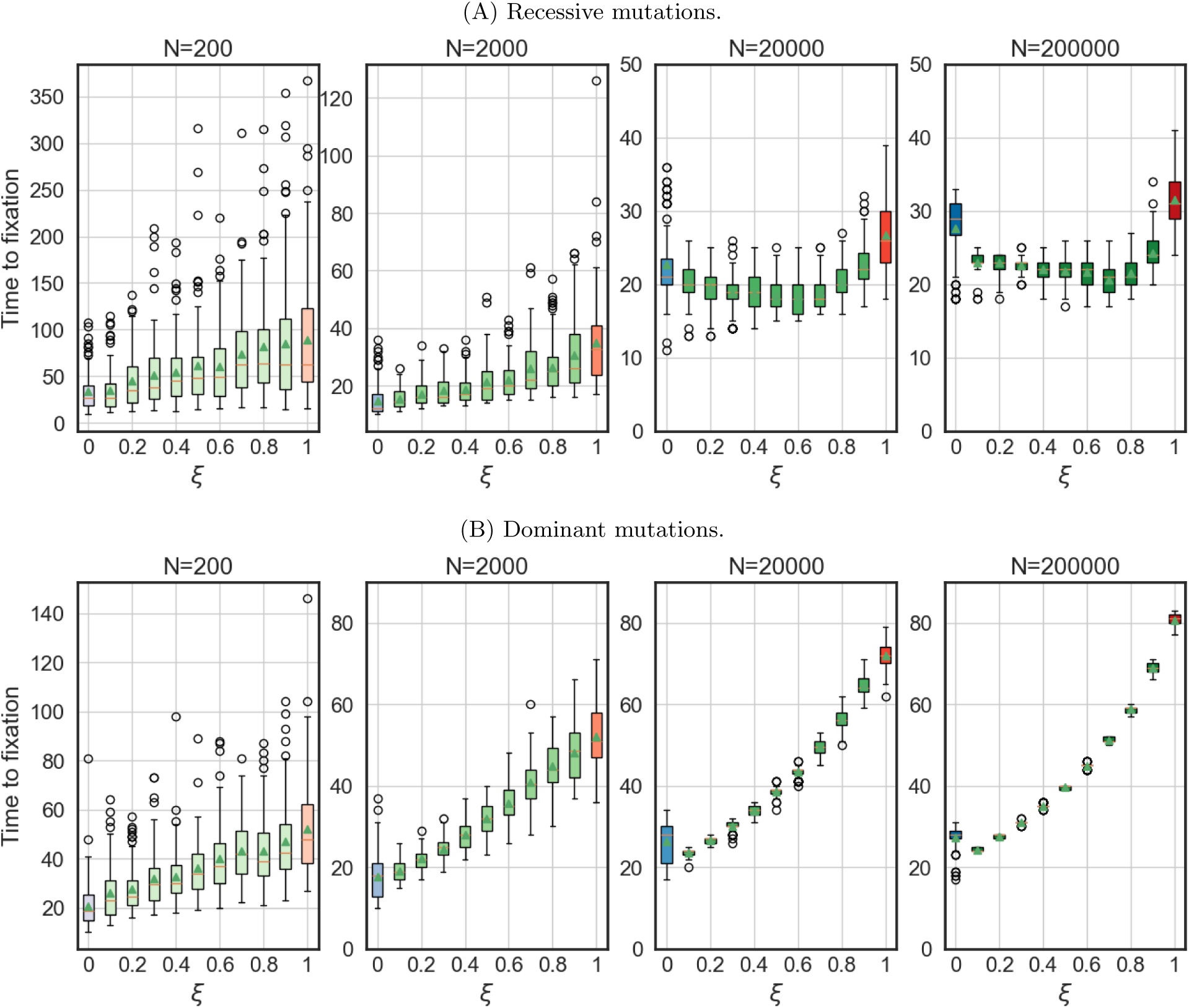
Time to fixation of the first beneficial allele for various fractions of mating hermaphrodites *ξ*. Resistance is encoded by six loci. Note the various scales on the Y-axis. Box plots show the median, interquartile range, and whiskers extending to 1.5x the interquartile range from the box edges.

Figure 5 demonstrates that while selfing results in the fastest fixation of the mutated allele in small populations, a combination of mating and selfing proves advantageous in larger populations. The proportion of mating that appears beneficial increases with population size.

Furthermore, this relationship is influenced by dominance. For equivalent population sizes, a considerably lower mating frequency led to the shortest fixation time when mutations are dominant. In populations of 20000 and 200000 individuals, the shortest time to fixation was observed under different conditions. When

mutations were dominant, the shortest time occurred when only 10% of hermaphrodites outcrossed. In contrast, when mutations were recessive, the shortest time occurred when 60% or 70% of hermaphrodites outcrossed, respectively. Again, this highlights the differences between *C. elegans*, that maintain males on relatively low frequencies, and dioecious parasitic worms, particularly when resistance mutations are dominant.

### 4.4 Dominant mutations lead to faster adaptation to higher drug concentration

Comparing Figure 3 and Supplementary Figure S2 demonstrates that adaptation to higher concentrations is faster when mutations are dominant. This is expected, as dominant mutations are exposed to selection even in the heterozygous state, conferring an immediate fitness advantage.

The time to fixation of the first mutation (Figure 5) is also generally longer for recessive mutations, especially in dioecious populations. This difference is less pronounced in androdioecious and hermaphrodite populations. We observed that the fastest adaptation always occurred for purely selfing populations or for populations with a high selfing rate. Figure 5 further reveals an unexpected interaction between population size and dominance. While recessive mutations require more time to fix in smaller populations, this is not the case for dominant mutations, where the differences between different population sizes were much smaller. This suggests that in smaller populations, mutations take longer to appear due to reduced mutation supply. However, once a beneficial mutation arises, it can reach prominence more rapidly due to stronger genetic drift in smaller populations. This effect is particularly pronounced for dominant mutations, which are immediately visible to selection, unlike recessive mutations that require homozygosity for phenotypic manifestation.

### 4.5 Mating, dominance and large population sizes hinder the fixation of mutations with small effects

As mentioned previously, dioecious populations exhibited limited fixation of recessive mutations, with an average of only three loci homozygous for the recessive allele by the simulation’s end. To further investigate this phenomenon, we analyzed the identity of the locus where the mutated allele first reached fixation (95% frequency). Supplementary Figures S3 - S4 reveal that in small, hermaphroditic populations, any of the six loci can be the first to reach fixation, suggesting a stochastic process driven by the random appearance of mutations. However, in larger populations, fixation consistently occurs first at one of the loci with a large effect. This indicates that in larger populations, the evolutionary trajectory is shaped primarily by competition between mutants, where those with larger effects have a selective advantage. Furthermore, both dominance and the presence of mating increased the likelihood of large-effect loci reaching fixation first, highlighting the influence of these factors on the adaptive process.

### 4.6 Mating increased the diversity

Mating is known to have significant impacts on genetic diversity, with selfing leading to much lower diversity than mating [Ingvarsson, 2002, Burgarella et al., 2024]. Diversity is also expected to be lower in smaller populations. Figure 6 depicts the diversity, calculated as Shannon index, over time (see Supplementary figure S5 for a zoomed-in version of the first 50 generations). In agreement with expectations, it shows that diversity is the lowest in the strictly selfing populations and smallest populations. However, it shows that in androdioecious populations, diversity is as high as in strictly mating populations. In simulations where only two loci encode variance, the diversity is even higher in androdioecious populations than in dioecious ones (Supplementary Figures S6 and S7). In all scenarios, we observed an initial steep increase in genetic diversity as new mutations appeared and increased in frequency. This was followed by a rapid decline as sensitive alleles were driven to extinction. The first peak in diversity can be attributed to the emergence and fixation of mutations with large effects. In contrast, the second peak arises from the increasing prevalence of mutations with smaller effects, which eventually also become fixed in the populations (except for dioecious ones). Since the selection advantage provided by these small-effect loci is weak, the adaptive process is slower, resulting in a broader peak. Notably, we did not observe a second peak in dioecious populations with recessive mutations, which aligns with our earlier finding that only three loci became fixed in this scenario.

**Figure 6:**
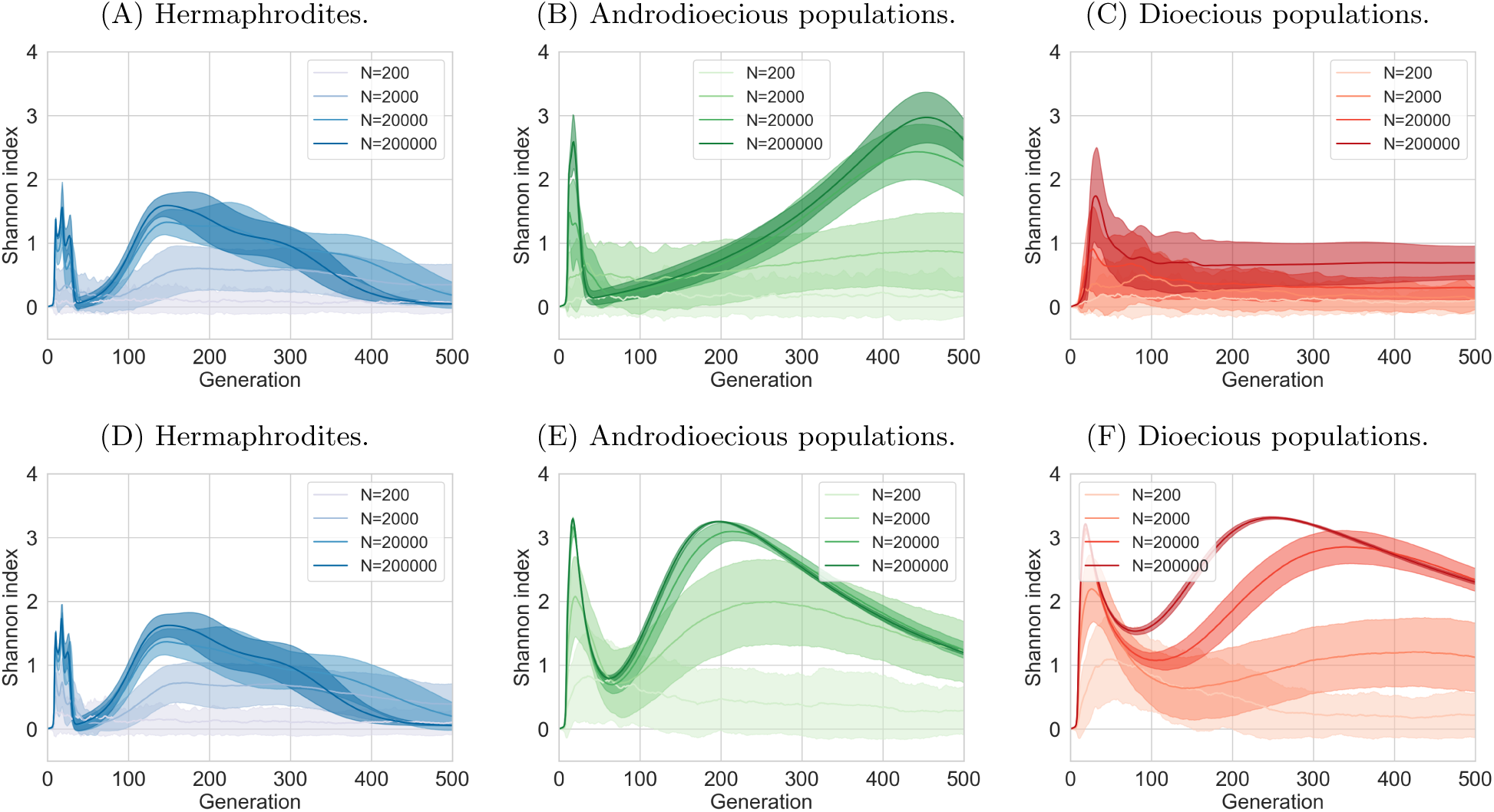
Biodiversity changes over time. Recessive and dominant mutations, six loci encoding resistance. A-C: recessive mutations, D-F) dominant mutations. In androdioecious populations, 50% of hermaphrodites reproduce by mating, 50% by selfing.

### 4.7 Short-term observations can contradict the long-term ones

Simulations were conducted to investigate population dynamics over 500 generations, equivalent to more than four years of continuous *in vitro* experimentation and at least 80 years of continued *in vivo* selection of parasitic nematodes. In addition to being time-consuming, such experiments are likely to be extremely costly and unlikely to be conducted often. In practice, experiments containing approximately 50 or fewer generations are more feasible [Hellinga et al., 2024]. However, our findings demonstrate that observations from shorter timeframes may yield conclusions that differ significantly from those derived from longer-term investigations.

Figure 7 shows the drug concentration that was reached in the experiments with populations with a range of mating systems (from hermaphroditism, *ξ* = 0, to dioecy, *ξ* = 1) after 10, 25, 100, 250 and 500 generations. ‘Drug concentration reached’ refers to a concentration at which the worms were able to survive in a particular generation but did not produce a sufficient eggcount (20 on average), allowing us to increase the drug concentration further. The adaptive advantage conferred by various mating systems varies considerably across different time points and population sizes (see Supplementary Figure S8 for bigger populations). Specifically, at early time points, hermaphroditic populations appear to exhibit the most rapid adaptation, with sexual reproduction seemingly conferring a disadvantage across all population sizes. This trend, however, reverses over time as the benefits of outcrossing become increasingly evident, particularly in populations with recessive mutations (see Supplementary Figure S9 for dominant mutations). These results highlight the critical importance of incorporating a temporal dimension into evolutionary studies, especially when considering the interplay between reproductive strategies and multi-locus evolution.

**Figure 7:**
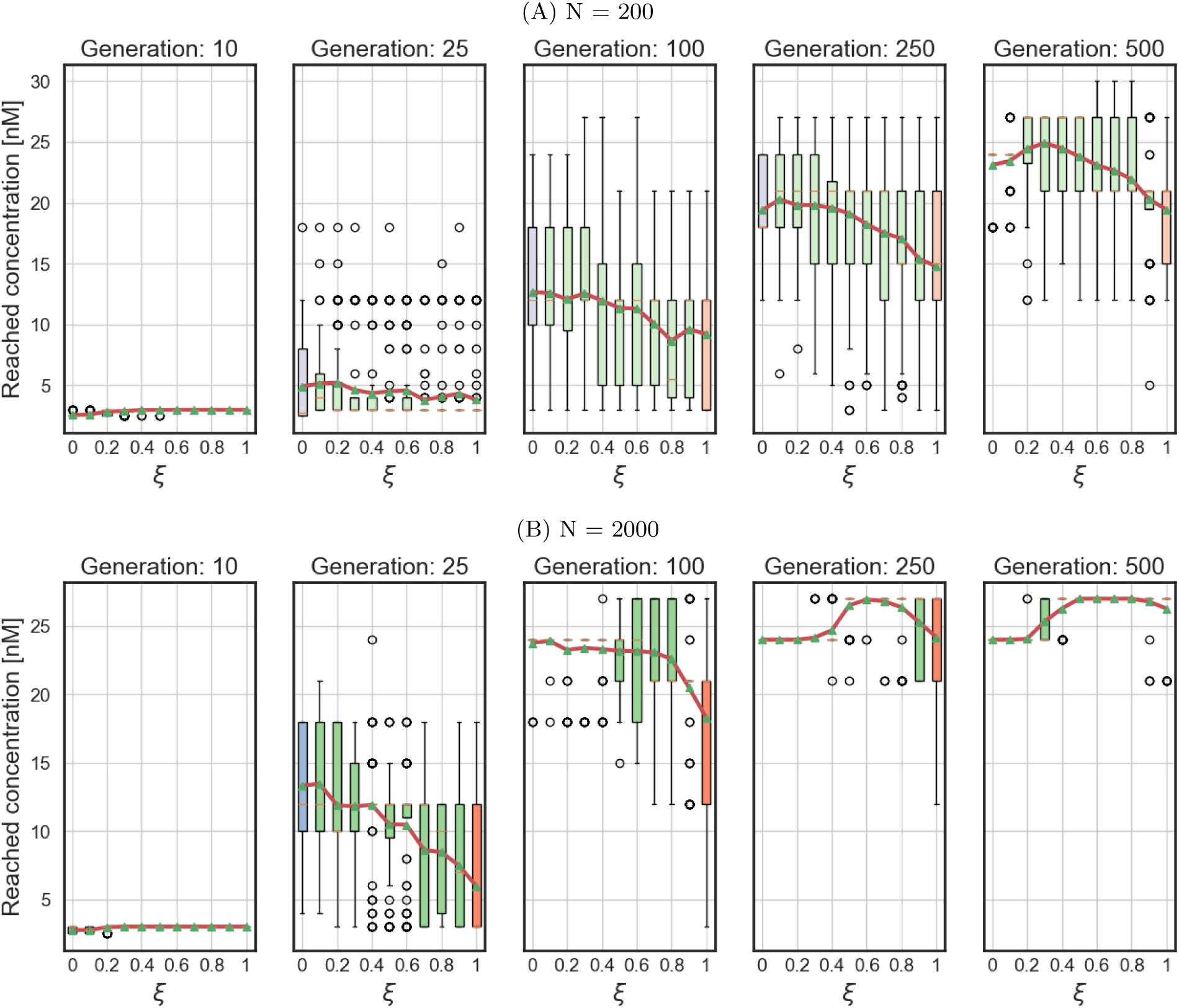
Drug concentration reached by populations at different time points, recessive mutations. Blue represents hermaphrodite populations, red dioecious ones, and green various levels of androdioecy. Box plots show the median, interquartile range, and whiskers extending to 1.5x the interquartile range from the box edges. Red line connects the means.

## 5 Discussion

Our study investigated how different mating systems influence the evolution of drug resistance in worm populations. We examined a range of mating systems, including dioecy, hermaphroditism, and androdioecy as an intermediate state. Through simulations based on the biology of *C. elegans* and experiments conducted over 500 generations, we discovered that mating systems significantly impact evolutionary trajectories.

Hermaphrodites, capable of self-fertilization, exhibited faster adaptation under recessive mutations. Selfing promotes homozygosity, effectively exposing and fixing new mutations [Ellis and Lin, 2014]. This is due to the increased probability of generating homozygous offspring from a single mutant individual, allowing for rapid fixation of the beneficial allele. In contrast, dioecious populations, reliant on outcrossing, faced a greater challenge in establishing recessive mutations, as these require the rare occurrence of mating between two carriers of the allele.

It is worth noting that the classic “cost of males” does not apply in our model, as the presence of males actually leads to increased average egg production per individual. This increase in fecundity with the mating fraction *ξ* arises from the fact that mated hermaphrodites lay more than thrice the number of eggs compared to those that reproduce through self-fertilization [Kimble and Ward, 1988]. Consequently, the average fecundity of a population—or metapopulation, when focusing on a specific strain—depends on the proportion of mated hermaphrodites. This reflects the maximum fecundity in the absence of drugs, where the actual egg count is influenced by the fitness of the strain, which is determined by both genetic and environmental factors.

This deviation from traditional theoretical models results in altered selective pressures whereby dioecious populations generate more offspring than hermaphrodites. This increased reproductive output enables dioecious populations to achieve higher drug concentrations despite initially adapting and accumulating mutations at a slower rate. Thus, over time, dioecious populations manage to survive and reproduce in environments with higher drug concentrations.

Androdioecious populations, characterized by a combination of selfing and outcrossing, appear to be able to access the advantages of both reproductive modes. Androdioecious populations adapted to increasing drug concentrations at a speed initially comparable to hermaphrodites and exceeded the speed of dioecious populations. While hermaphrodites lagged in their ability to reach higher concentrations, androdioecious populations, after a delay, successfully adapted to it. This pattern suggests that the flexibility afforded by androdioecy allows populations to rapidly exploit beneficial mutations through selfing while maintaining sufficient genetic diversity through outcrossing to overcome more long-term selective pressures. These results align with previous empirical observations of lower extinction rates in androdioecious lineages of *C. elegans* during experimental evolution in a new environment, supporting the adaptive benefits of combining selfing and outcrossing [Morran et al., 2011, Masri et al., 2013, Chelo and Teotónio, 2013].

Our study allowed us to investigate a range of reproductive strategies, ranging from pure selfing to pure mating, with intermediate stages. We showed that the optimal balance between mating and selfing depends both on genetics - whether the mutations are dominant or recessive, as well as on the population size. A small fraction of males and mating may lead to the fastest adaptation. This is consistent with three independent studies that identified an advantage of outcrossing of androdioecious *C. elegans*populations during coevolution with pathogenic bacteria. Two of these studies included coevolving *Bacillus thuringiensis* as an antagonist and demonstrated that the favoured outcrossing rate never reached 100%, but remained at an intermediate level of 20-60% [Masri et al., 2013, Schulte et al., 2010]. In these cases, outcrossing with males was directly shown to increase offspring resistance against the pathogen [Masri et al., 2013]. The third coevolution experiment, which included a *C. elegans* population with less genetic diversity than in the above studies, and also a different pathogen, *Serratia marcescens*, found a steady increase in outcrossing rates from 20% to above 70% [Morran et al., 2011]. Overall, these findings strongly suggest an advantage of the intermediate outcrossing rates facilitated by the androdioecious reproductive system.

Our findings further highlight the role of population size in shaping adaptive outcomes. We observed that in small populations, any locus could be the first to reach fixation, indicating a stochastic process driven by the chance emergence of beneficial alleles. Conversely, in larger populations, adaptation becomes more predictable, with loci harbouring large-effect mutations consistently reaching fixation first. This shift suggests that competition between mutants, rather than mere chance, dictates the evolutionary trajectory in larger populations. Furthermore, if natural selection favours the recombination of advantageous alleles at two loci or if it favours heterozygosity (not investigated in this model), then adaptation should be faster in large populations. One example of such a scenario comes from previous evolution experiments, in which *C. elegans* coevolved with its pathogen *B. thuringiensis* and where reciprocal co-adaptations were favored in larger host populations [Papkou et al., 2021].

Most importantly, our study reveals the strong influence of that mating strategy on evolutionary dynamics and its complex interaction with other factors, such as population size and genetic structure. This has important implications for the use of *C. elegans* as a model species for the study of drug resistance in parasitic nematodes that are, to our knowledge, almost entirely dioecious. It suggests that androdioecious *C. elegans* benefits from the high selfing rates that lead to fast adaptation, especially in relatively short timescales typically used in evolutionary experiments. By generalizing the finding for *C. elegans* to other species, we may overestimate their ability to evolve resistance and, in turn, suggest non-optimal drug treatments with possible detrimental effects. At the same time, our findings indicate that androdiecious pathogenic helminths, should they emerge, will have a higher risk of evolving resistance against anthelmintics.

Our modelling study further highlights the necessity of theoretical approaches complementary to experimental observations, such as our model, in order to capture the full complexity of evolutionary dynamics. Mathematical models and simulations are especially valuable when conducting laboratory experiments that are impractical or unfeasible, such as when long time-scales, large population sizes or high replication numbers are needed. In particular, our long-term simulations underscore the limitations of empirical studies when investigating the evolutionary processes across long time periods. While practical constraints often limit *in vitro* and *in vivo* experiments to a feasible timeframe of 50 or fewer generations, our model reveals that different outcomes can be observed over periods of many hundreds of generations. This is particularly evident in the context of reproductive mode, where the apparent short-term advantage of hermaphroditism can give way to the long-term benefits of outcrossing.

It is important to acknowledge certain limitations of our model. We assumed unlinked autosomal genes, which may not fully reflect the complexities of real-world genomes and considering the linkage between the genes is an important expansion of our model. Furthermore, we assumed that both the benefits and costs of individual mutations combine in an additive manner. While this is a common assumption in many polygenic models, several recent studies suggest that mutations may interact in a multiplicative way instead. Using a multiplicative approach can be beneficial, especially for cost, because it prevents negative fitness values, even when multiple costly mutations are combined, thus allowing for the simulation of such mutations. However, we used an additive model to avoid introducing epistasis into our analysis. Epistasis occurs when the effect of one mutation depends on the presence of another, which would complicate the interpretation of our results and obscure the direct effects of the mutations themselves. While we recognize the significance of epistasis in evolutionary processes, our primary goal for this research was to concentrate on other effects, and the consideration of epistasis is a promising extension for future research.

The influence of drug concentration on egg count was modelled solely through hermaphrodites, neglecting any potential role of male fitness. More specifically, we simplified the impact of anthelmintics on worm fitness by focusing solely on reduced egg count, regardless of the underlying causes—such as a shorter lifespan of the hermaphrodite, lower fecundity, or decreased likelihood of eggs hatching. We did not consider other factors, like a reduced chance of finding a mating partner. While this simplification was necessary, the explicit consideration of a larger number of factors represents a worthwhile avenue for future work. In particular, future studies could address these limitations by incorporating linked genes, exploring the impact of male fitness on adaptation, and extending the model to encompass other reproductive modes, such as asexual reproduction observed in some parasitic worm species.

Finally, we note that the gradually increasing drug concentrations modelled here are more representative of the laboratory experiments than of the anthelmintic use in real populations. However, as our main objective was to discuss the suitability of *C. elegans* for studies of drug resistance evolution in the laboratory setting, we chose to gradually increase drug concentration as it agrees well with methods employed by most researchers in order to select resistant parasite strains in animal experiments. Moreover, we believe that this study is also relevant for natural populations. Pathogens are often exposed to lower effective concentrations than those that would kill them. For instance, reinfection can occur when drug concentrations have already dropped to levels that do not completely prevent further parasite development but still select for certain parasite genotypes. In addition, drug concentration is not constant throughout the human and animal body, and some worms may experience higher concentrations, while some lower. The differences also occur during the course of drug metabolism and between hosts who differ in their metabolism and uptake of the drugs.

In conclusion, our study demonstrates the complex interplay between mating systems, population size, and genetic architecture in driving the evolution of drug resistance in worms. While hermaphroditism initially facilitates rapid adaptation through selfing, dioecy and androdioecy ultimately prevail in high drug concentrations due to increased genetic diversity and offspring production. Notably, androdioecy, by combining selfing and outcrossing, appears to offer the most robust adaptive strategy. These findings underscore the importance of long-term evolutionary perspectives and integrated theoretical approaches for understanding and predicting drug resistance dynamics in parasitic worms, with implications for effective control strategies. Our model, in combination with the experimental system, could be used for the testing of new drugs and the general potential of nematodes to evolve resistance under a variety of conditions. This approach should be expanded in the future to include nematodes with different reproductive strategies, especially those of parasitic worms, and population structures that permit migration and gene flow, reflecting natural complexity.

## Acknowledgements

This work has been supported by Swiss National Science Foundation. We thank Maya Louage for proofreading and correcting the simulation code, as well as two anonymous reviewers for their constructive comments on the previous version of the manuscript.

## Supplementary figures

**Figure S1:**
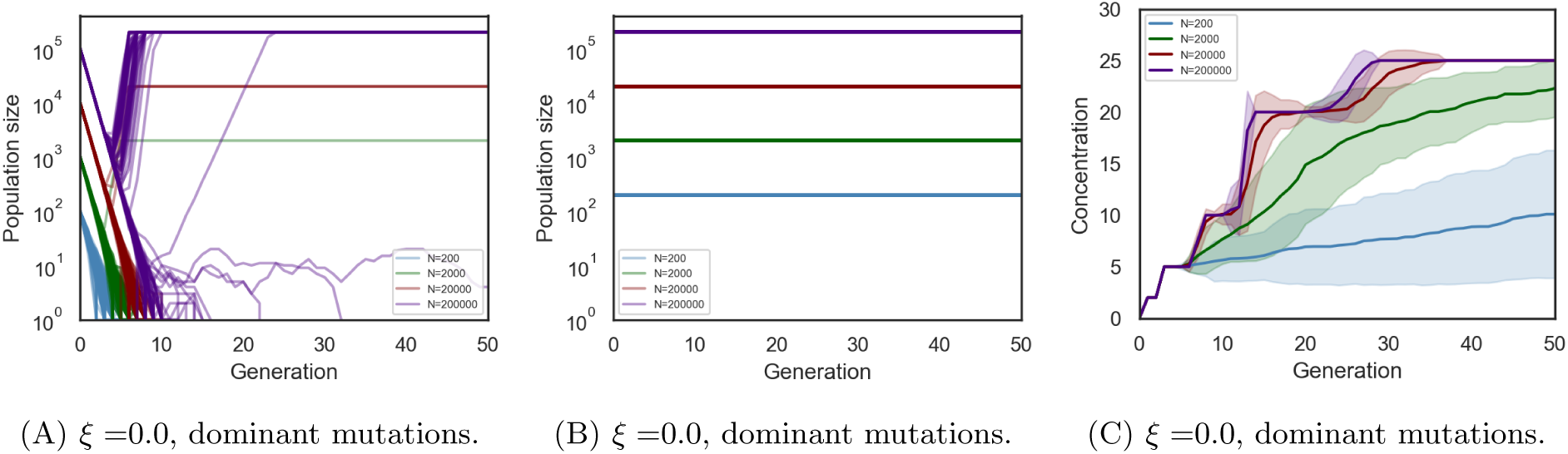
Sudden vs gradual drug exposure. A) Worm populations exposed to higher drug concentrations (10 nM) without prior adaptation to lower concentrations are likely to go extinct, especially if the population sizes are small. B) Populations exposed to gradually increasing concentration, as shown in (C), survive and adapt. (C) Surviving and expanding populations allow for a gradual increase in concentration every few generations.

**Figure S2:**
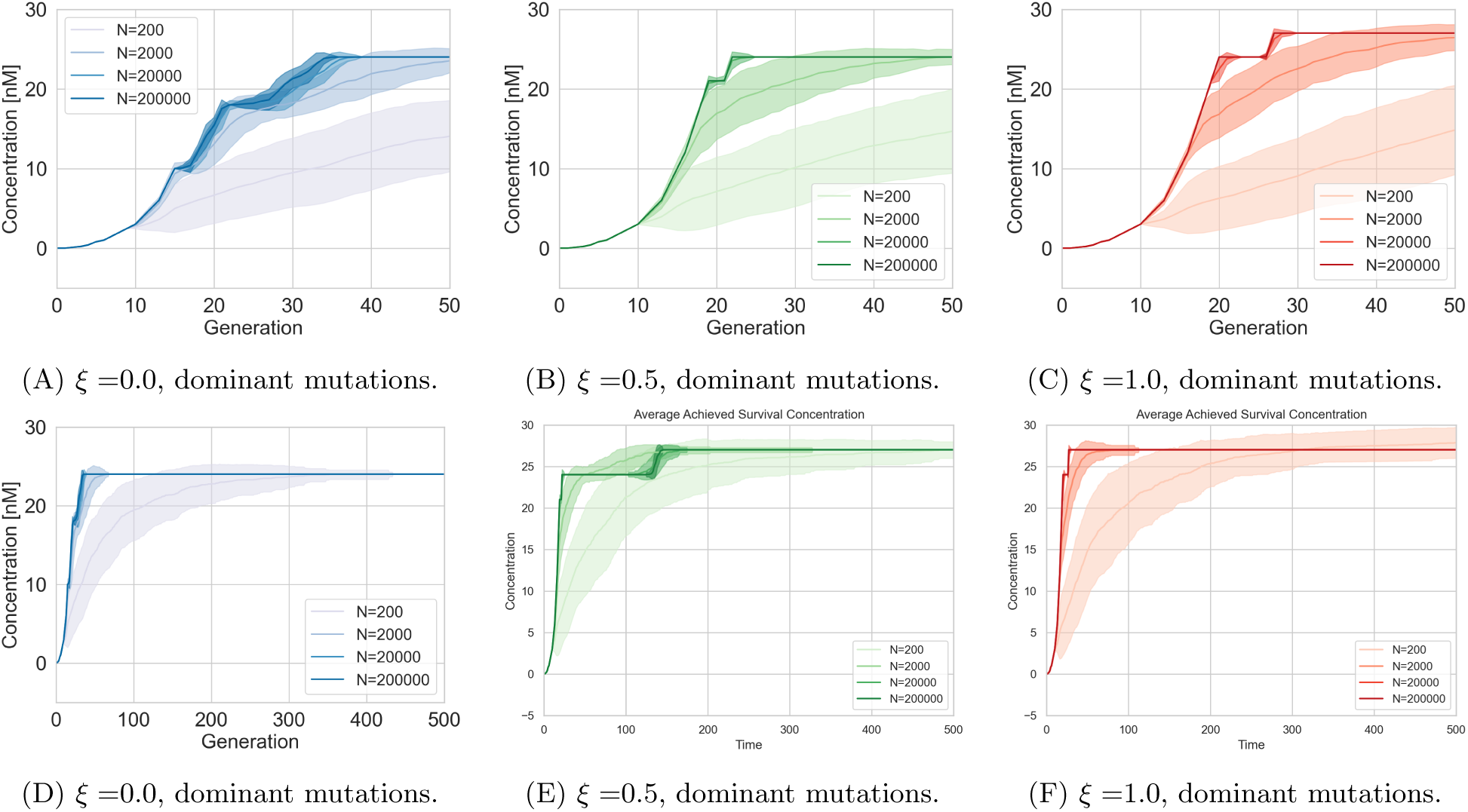
Adaptation in increasing drug concentration for dominant mutations. A-C) first 50 generations, D-F) all 500 generations. Mean and standard deviation of 100 simulation trials.

**Figure S3:**
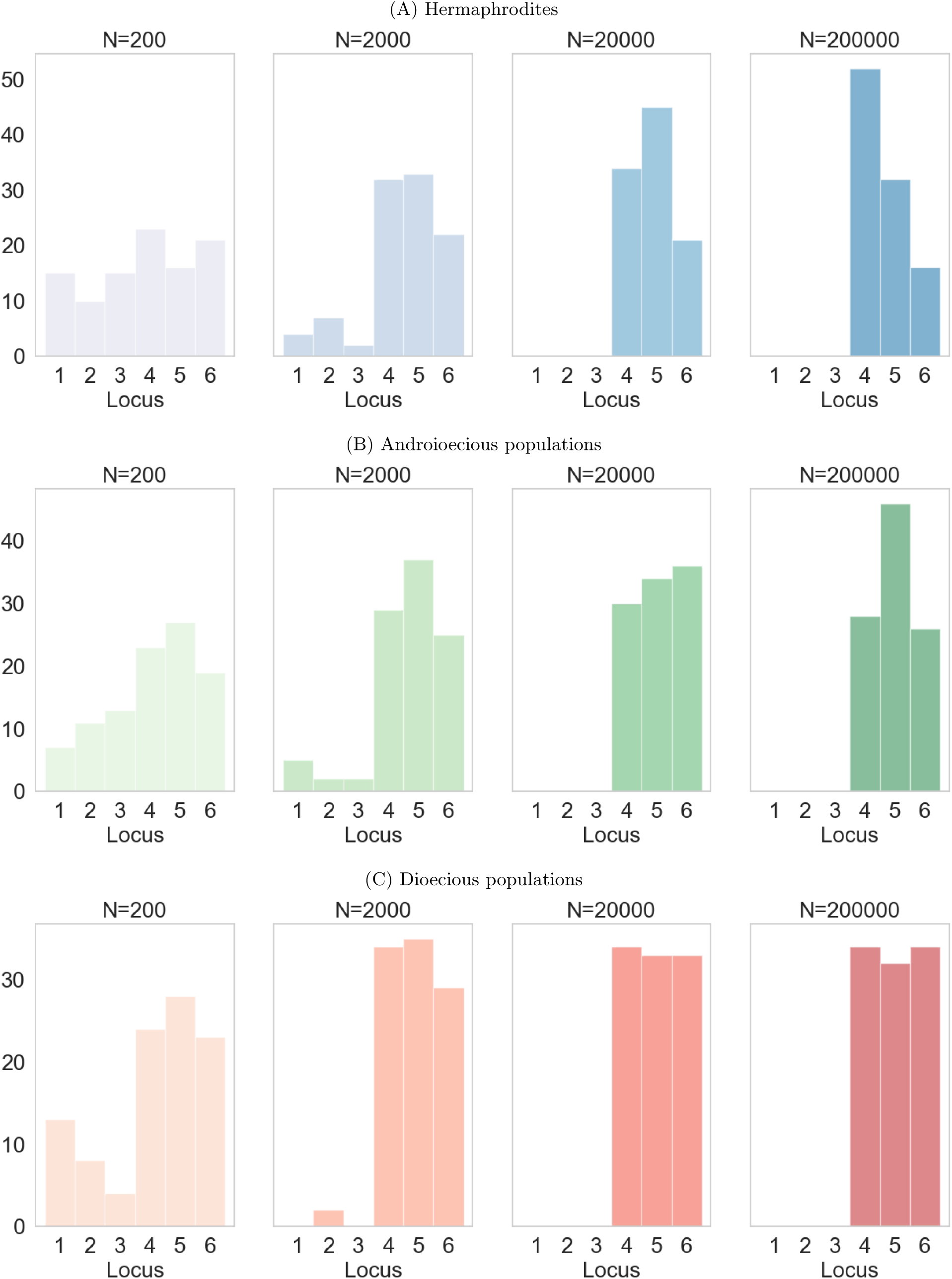
Time to fixation of the first mutation, recessive mutations. Note that there are different axes.

**Figure S4:**
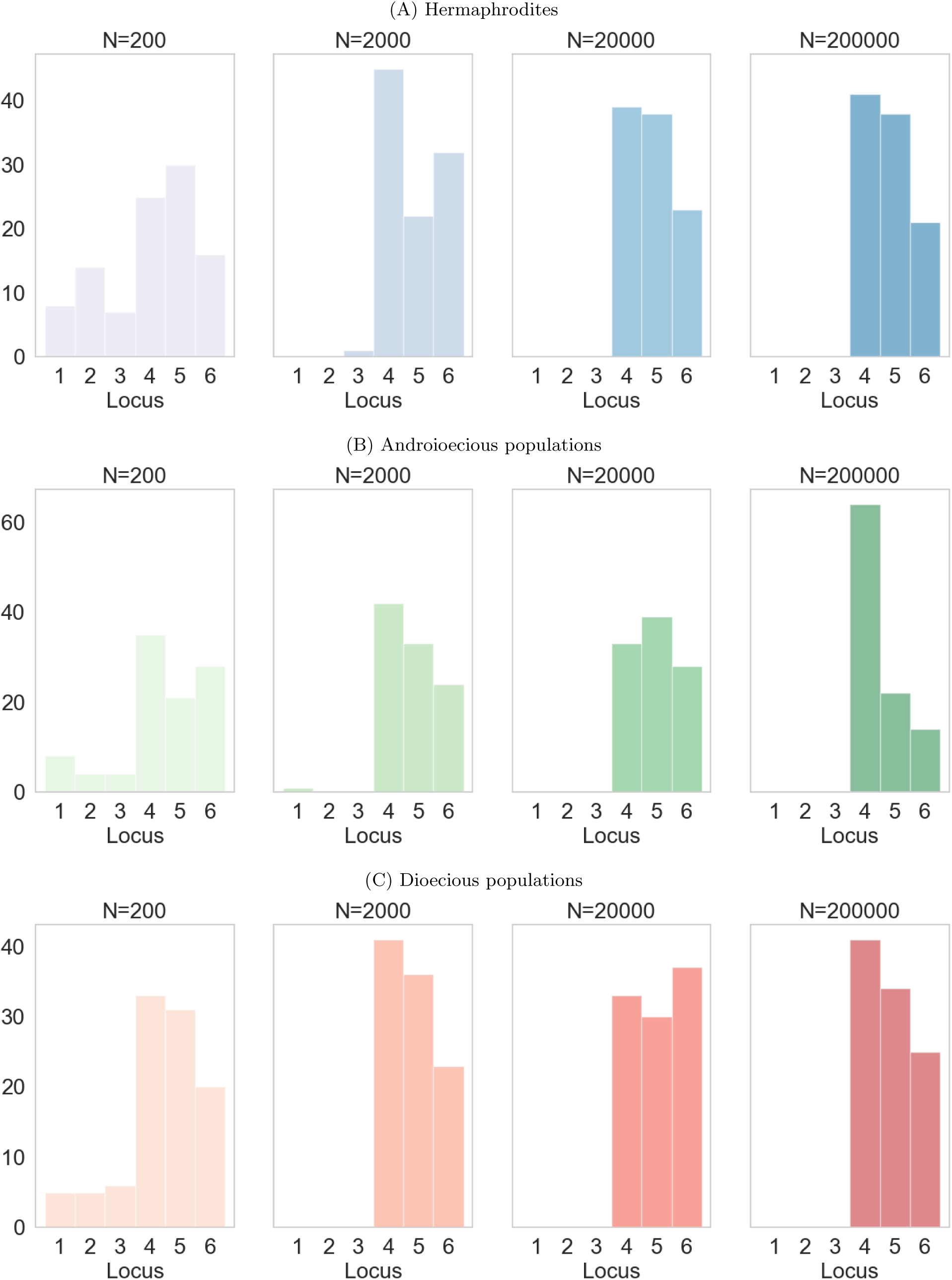
Time to fixation of the first mutation, dominant mutations. Note that there are different axes.

**Figure S5:**
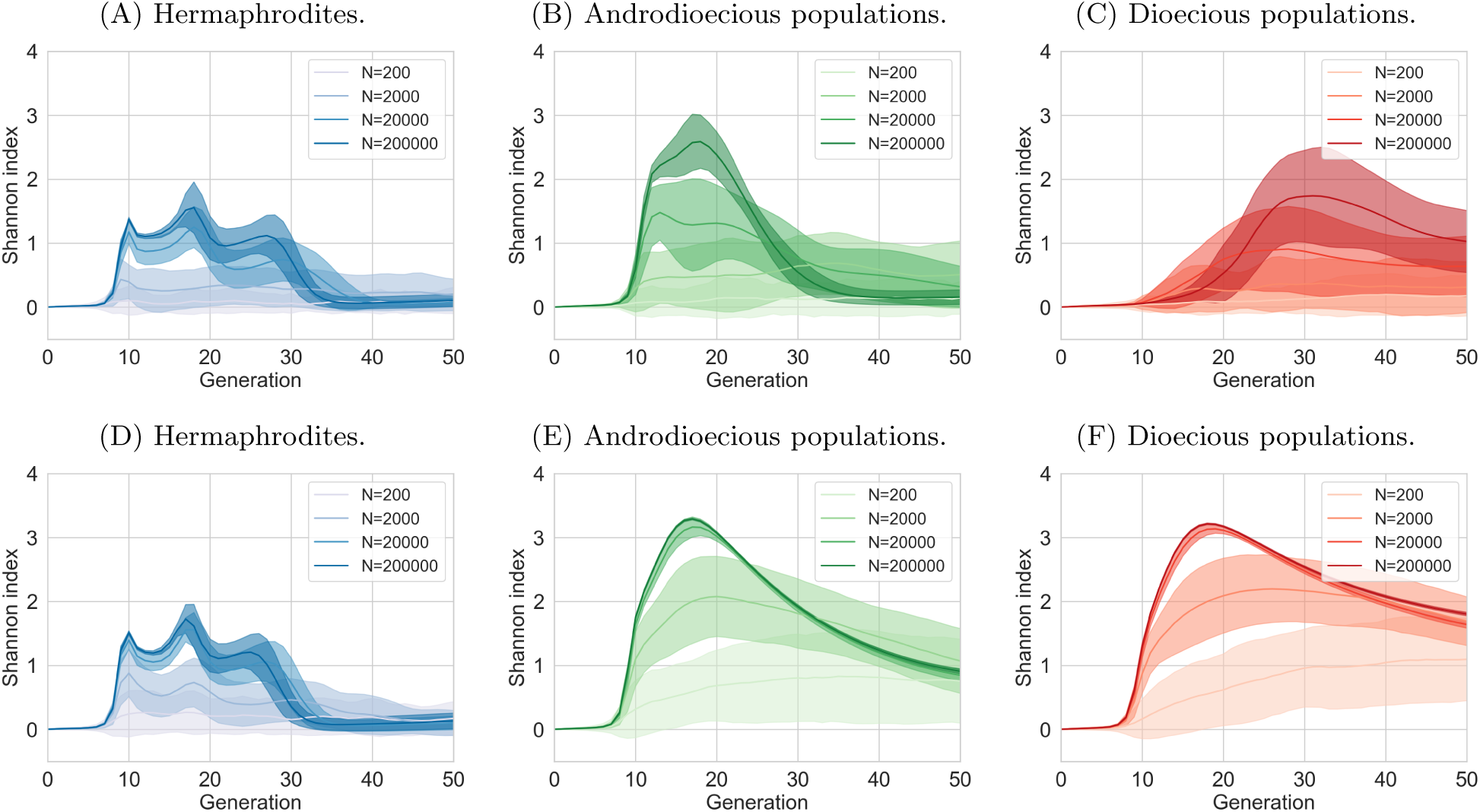
Biodiversity changes over time. Recessive and dominant mutations, six loci encoding resistance. A-C: recessive mutations, D-F) dominant mutations. In androdioecious populations, 50% of hermaphrodites reproduce by mating, 50% by selfing. First 50 generations.

**Figure S6:**
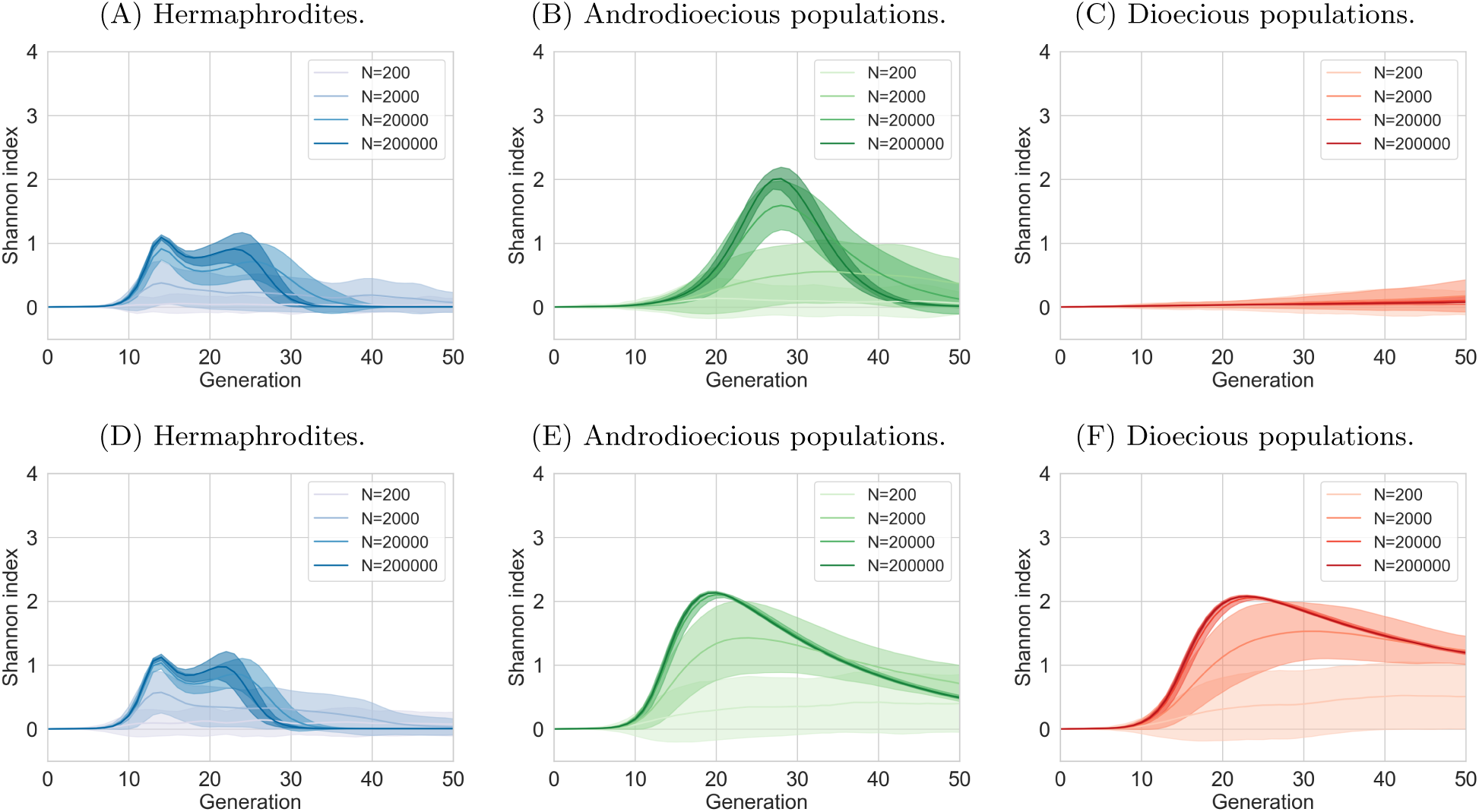
Biodiversity changes over time. Recessive and dominant mutations, two loci encoding resistance. A-C: recessive mutations, D-F) dominant mutations. In androdioecious populations, 50% of hermaphrodites reproduce by mating, 50% by selfing. First 50 generations.

**Figure S7:**
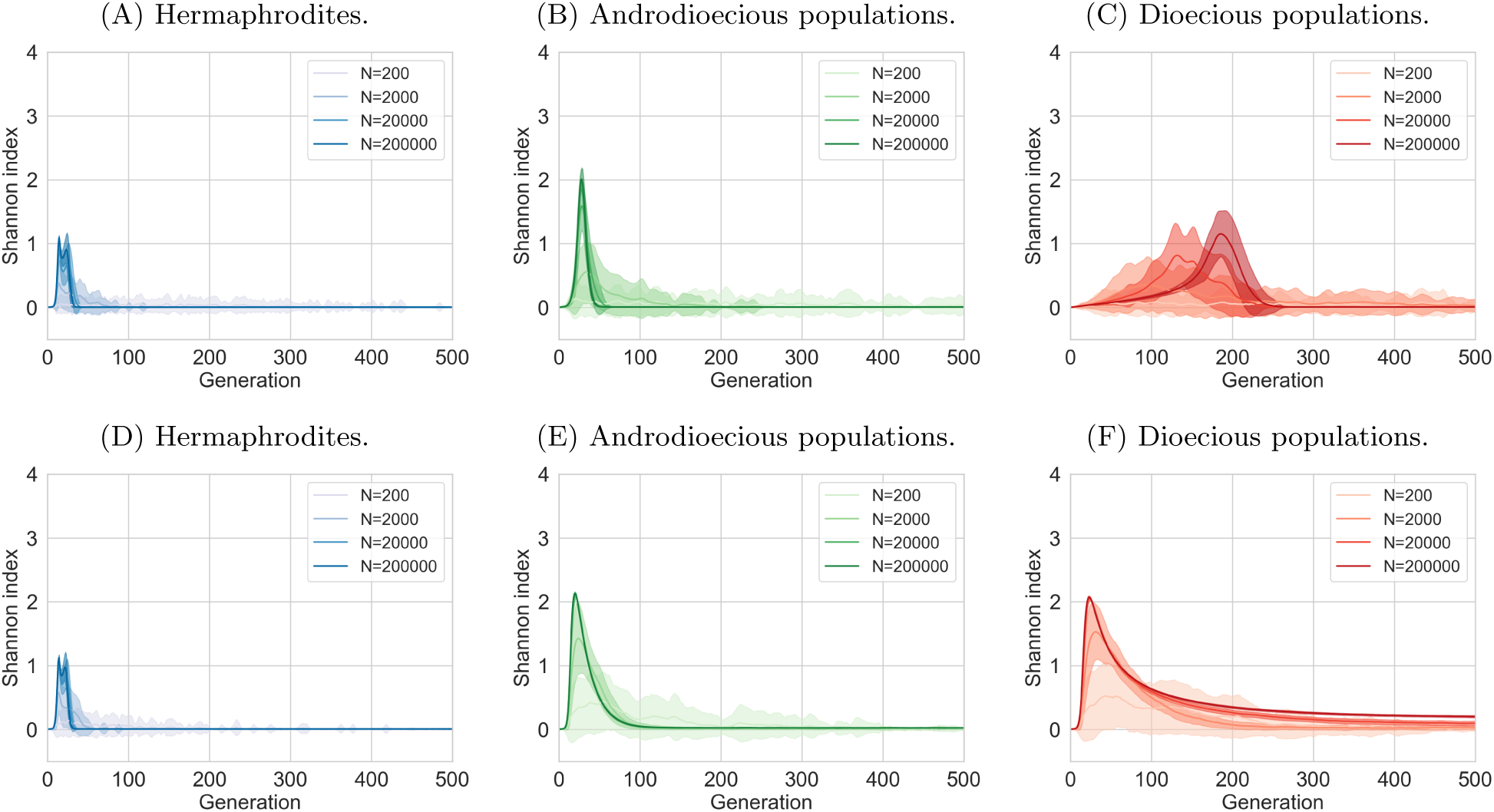
Biodiversity changes over time. Recessive and dominant mutations, two loci encoding resistance. A-C: recessive mutations, D-F) dominant mutations. In androdioecious populations, 50% of hermaphrodites reproduce by mating, 50% by selfing. All 500 generations.

**Figure S8:**
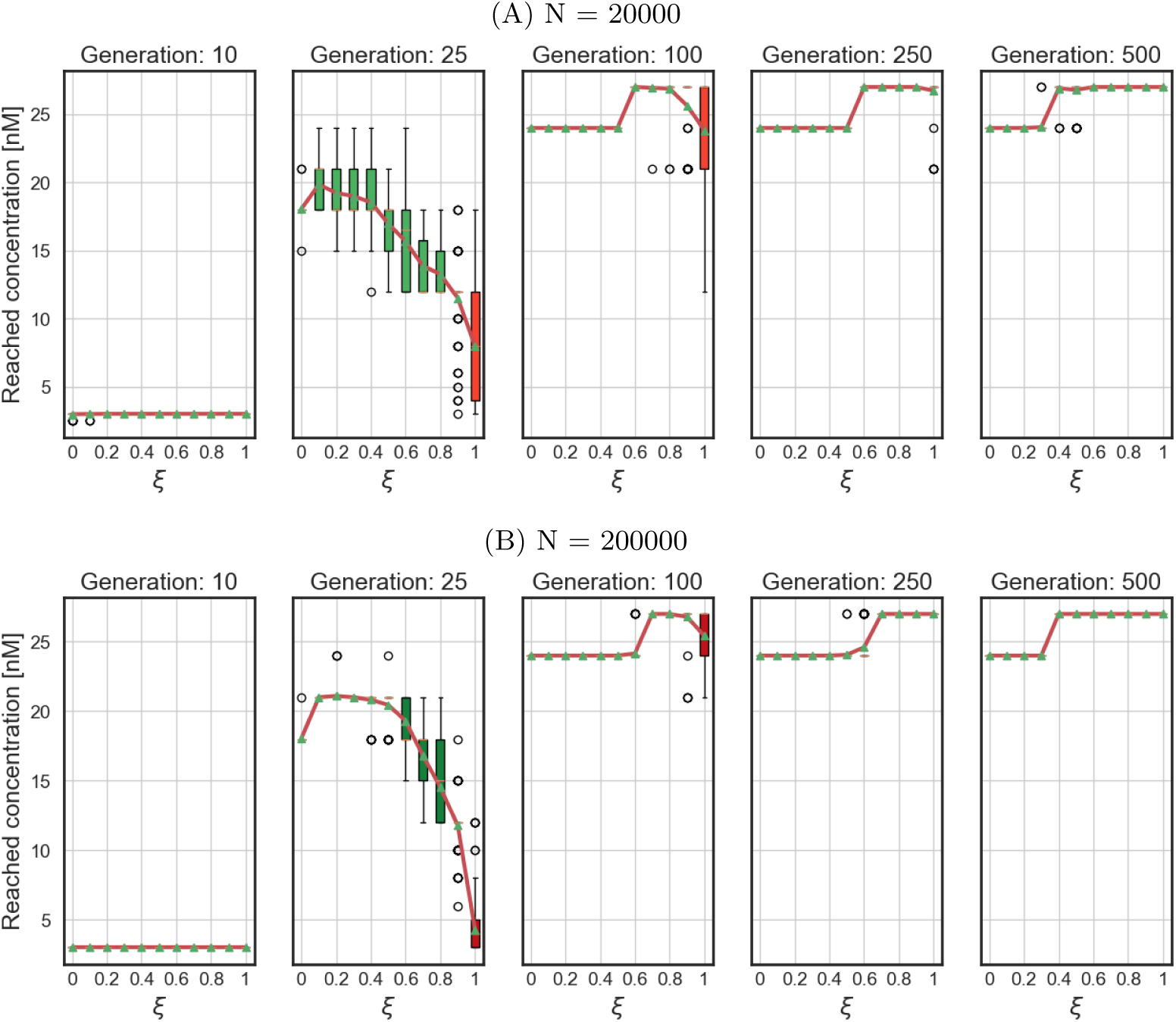
Drug concentration reached by populations at different time points, recessive mutations, large populations.

**Figure S9:**
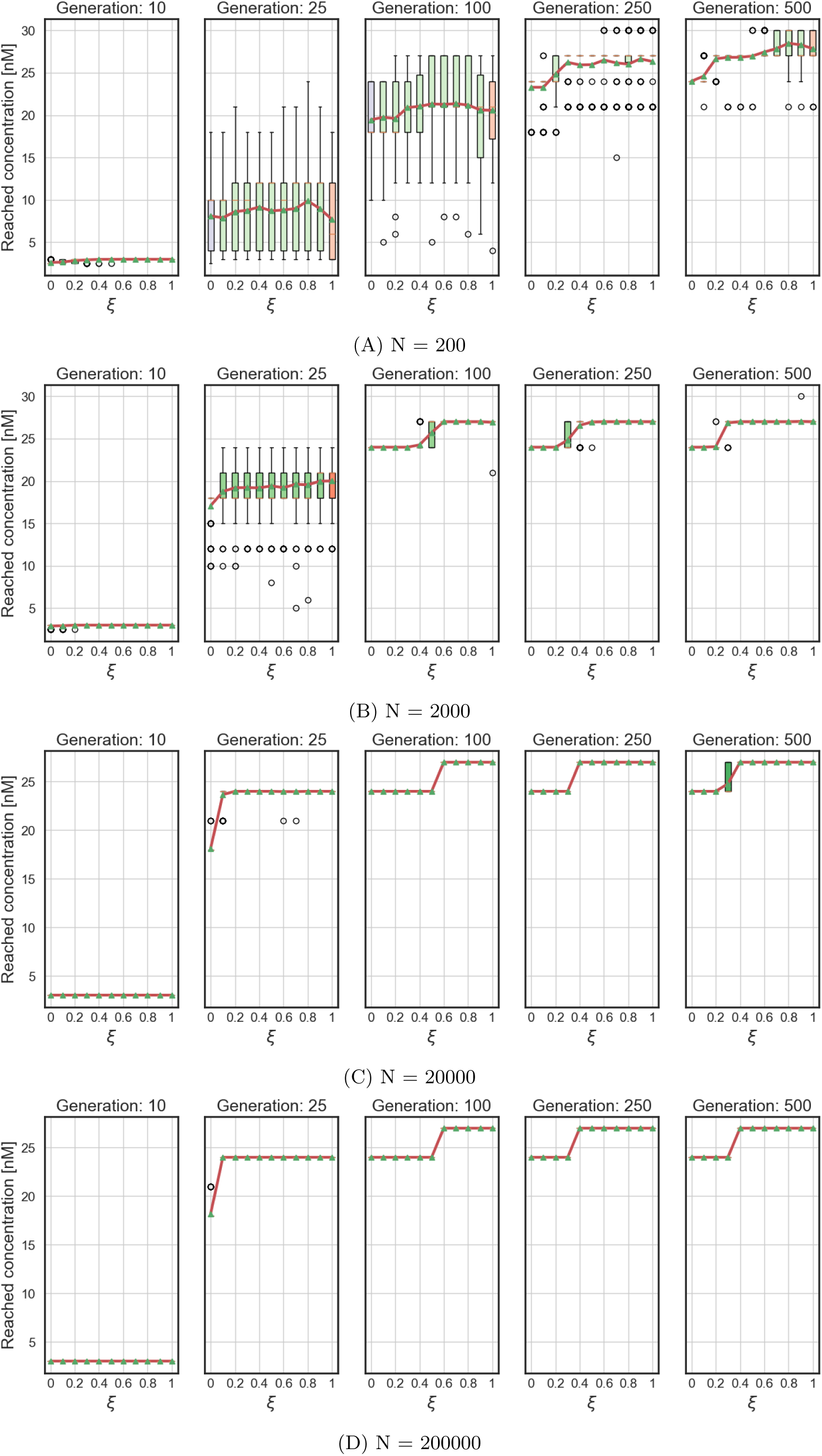
Drug concentration reached by populations at different time points, dominant mutations.

